# Single-Domain Antibody-Based Protein Degrader for Synucleinopathies

**DOI:** 10.1101/2024.03.11.584473

**Authors:** Yixiang Jiang, Yan Lin, Amber M. Tetlow, Ruimin Pan, Changyi Ji, Xiang-Peng Kong, Erin E. Congdon, Einar M. Sigurdsson

## Abstract

Synucleinopathies are a group of neurodegenerative diseases characterized by the accumulation of α-synuclein (α-syn) in the brain, leading to motor and neuropsychiatric symptoms. Currently, there are no known cures for synucleinopathies, and treatments mainly focus on symptom management. In this study, we developed a single-domain antibody (sdAb)-based protein degrader with features designed to enhance proteasomal degradation of α-syn. This sdAb derivative targets both α-syn and Cereblon (CRBN), a substrate-receptor for the E3-ubiquitin ligase CRL4^CRBN^, and thereby induces α-syn ubiquitination and proteasomal degradation. Our results indicate that this therapeutic candidate enhances proteasomal degradation of α-syn, in addition to the endogenous lysosomal degradation machinery. By promoting proteasomal degradation of α-syn, we improved clearance of α-syn in primary culture and mouse models of synucleinopathy. These findings indicate that our sdAb-based protein degrader is a promising therapeutic candidate for synucleinopathies. Considering that only a small percentage of antibodies enter the brain, more potent sdAbs with greater brain entry than whole antibodies could enhance clinical benefits of antibody-based therapies.

## Background

Synucleinopathies, including Parkinson’s disease (PD), Lewy body dementia (LBD), and Multiple system atrophy (MSA), are a group of neurodegenerative diseases marked by aggregation of α-synuclein (α-syn) into Lewy bodies primarily within the central but to some extent within the peripheral nervous system (*1*). These diseases pose a considerable burden on caregivers, families, and society (*2*). Regrettably, no cure is presently available and therapeutic options mainly focus on managing the symptoms.

Familial forms of synucleinopathies are linked to mutations and replication of the SNCA (α-syn) gene, indicating that α-syn is the driving force for these diseases (*3*). Therefore, reducing α-syn accumulation is a key therapeutic strategy (*1, 3*). Different approaches are being pursued, including: 1) Active and passive immunization to promote clearance of α-syn aggregates (*4*); 2) siRNAs to decrease α-syn synthesis (*5, 6*); 3) boosting lysosomal and/or autophagic activity to increase α-syn degradation (*7–13*), and; 4) small molecule modulators to reduce α-syn aggregation (*14*). However, each approach has potential drawbacks. For example, immunization may lead to immunological side effects (*4*), siRNAs may cause unintended gene silencing (*15*), boosting lysosomal activity may have limited effectiveness due to pre-existing impairments and a lack of specificity that could disrupt normal cellular processes (*16, 17*), and small molecule inhibitors of protein aggregation may affect other proteins (*18*).

There is growing evidence to suggest that pathological α-syn can spread from cell to cell, leading to the spatiotemporal spread of pathologies associated with disease phenotypes (*19–24*). To counter this, and to target α-syn inside cells as well, antibody-based drugs that prevent the aggregation and spread of toxic α-syn may alter the course of disease (*25*). However, current clinical α-syn immunotherapies based on whole IgG antibodies (150 kDa) have difficulty crossing the blood-brain barrier, limiting their effectiveness (*26*). Notably, no major side effects have been observed in these trials alleviating the potential drawbacks noted above. To improve efficacy, one strategy is to use smaller antibody fragments, such as single-chain variable fragments (scFv, 25 kDa) or single-domain antibodies (sdAbs or VHHs, ∼15 kDa), which have greater brain uptake albeit shorter half-life than whole antibodies (*27, 28*).

A few studies have focused on the development of antibody fragments targeting α-syn, including both scFvs (*29–31*) and sdAbs (*32–35*). These fragments have shown promise in inhibiting α-syn assembly or/and preventing cellular toxicity, as demonstrated in in vitro studies. A recent investigation utilized synthetic nanobody libraries in yeast to develop an anti-α-syn sdAb, PFFNB2 (*36*). Intraventricular injection of adeno-associated virus (AAV) encoding PFFNB2 has been shown to impede the spreading of α-syn pathology in mice. However, its efficacy on α-syn degradation has not been reported.

Reliable markers to assess α-syn proteopathic burden and to identify the stage of pathology in patients remain a major obstacle for development of therapies. To address this, we recently developed sdAb-based in vivo imaging probes (2D10 and 2D8) which allow for specific and non-invasive imaging of α-syn pathology in mice, with the brain signal strongly correlating with lesion burden (*37*). These sdAbs were derived from phage display libraries developed from a llama immunized with full-length recombinant (rec) α-syn. They readily interact with pathological α-syn derived from both human and mouse brain samples, supporting their potential as therapeutic and diagnostic candidates.

We and others have shown that antibodies can target proteinopathies both intra- and extracellularly (*27, 38–45*). Antibodies that can enter neurons have the potential to bind to protein aggregates within the endosomal-lysosomal system and/or ubiquitin-proteasome system and promote their clearance. As α-syn is predominantly intracellular, targeting it within cells would be more effective than extracellular clearance alone. However, the activities of the lysosome and proteasome, the two primary intracellular protein degradation compartments, can be compromised by neurodegeneration and aging (*16, 46–48*). Malfunction of these organelles may exacerbate α-syn toxicity by accelerating its accumulation due to defective protein degradation.

Recently, PROteolysis TArgeting Chimeras (PROTACs), which are bifunctional compounds consisting of a ligand targeting a protein of interest (POI) and an E3 ligase ligand, have emerged as a promising therapeutic approach in drug discovery. They bring POI and E3 ligase into proximity, triggering proteasomal degradation (*49, 50*). To enhance the therapeutic potential of our anti-α-syn sdAbs, we conjugated the sdAbs with thalidomide, an E3 ligase ligand, to merge sdAb-based immunotherapy with PROTAC technology. In this study, we show that an sdAb-based protein degrader, 2D8-PEG4-Thalidomide (2D8-PEG4-T), enhances α-syn clearance in both primary culture and mouse synucleinopathy models by promoting its proteasomal degradation. These findings indicate that our sdAb-based protein degrader is a promising therapeutic candidate for synucleinopathies. Given that only a small percentage of whole antibodies enter the brain, more potent sdAbs with better brain penetration could enhance clinical benefits of antibody-based therapeutic candidates.

## Materials and Methods

### Mammalian expression and purification of single domain antibodies

The sdAb clones were expressed in a mammalian system to avoid having to endotoxin purify clones expressed in a bacterial system. pVRC8400-sdAb constructs were made by inserting their individual gene sequences between the 5’ EcoRI and 3’ AfeI sites with a signal peptide, mouse interleukin-2 (IL-2) leader sequence (MYRMQLLSCIALSLALVT) at N-terminus and myc-his tags at C-terminus for detection and purification. The sdAbs were then expressed in FreestyleTM 293F cells (Invitrogen, Cat. No. R790-07). Briefly, FreestyleTM 293F cells were transiently transfected with the mixture of DNA plasmid and polycation polyethylenimine (PEI; 25 kDa linear PEI, Polysciences, Inc., cat. No. 23966). The transfected cells were incubated at 125 rpm in FreestyleTM 293 Expression Medium for suspension culture at 37°C with 5% CO_2_. The supernatants were harvested 5 days after transfection, filtered through a 0.45 μm filter, followed by sdAb purification using Ni-NTA columns (GE Healthcare). The wash buffer consisted of 20 mM Na_3_PO_4_, 100 mM NaCl, and 20 mM imidazole, pH 7.4. The elution buffer consisted of 20 mM Na_3_PO_4_, 100 mM NaCl, and 500 mM imidazole, pH 7.4. Following elution, the sdAb was dialyzed into PBS and its concentration determined by a BCA assay.

### Synthesis of single domain antibodies conjugate

To prepare the thalidomide-O-(PEG)n-modified sdAb 2D8, a 1 mL solution of 1 mg/mL sdAb 2D8 was prepared in 1X PBS (pH 7.4). Prior to use, a 50 mM solution of thalidomide-O-(PEG)n-NHS ester (n=2, 4, 6) was freshly prepared in DMSO. Ten equivalents of the thalidomide-O-(PEG)n-NHS esters were added to the sdAb solution, followed by thorough mixing. The mixture was then incubated at room temperature for 1 h, after which any unreacted thalidomide-O-(PEG)n-NHS ester was removed by dialysis.

### Bio-layer interferometry binding affinity assay

The bio-layer interferometry (BLI) experiments were carried out using a ForteBio Octet® RED96 instrument in a 96-well black flat bottom plate with shaking speed set at 1,000 rpm, at room temperature. Samples were prepared in 200 μL of 1X PBS assay buffer containing 0.05% Tween-20 at pH 7.4.

To measure the biophysical interaction between the his-tagged sdAbs and the different α-syn preparations (rec α-syn and human LBD brain-derived soluble α-syn S1 fraction), we used the Ni-NTA biosensor, which is pre-immobilized with nickel-charged tris-nitrilotriacetic (Tris-NTA) groups to capture his-tagged molecules. Before the BLI analysis, the sensor was hydrated in the assay buffer for 10 min, and a baseline was established in the buffer for 120 sec. The Ni-NTA tips were then loaded with his-tagged sdAbs at 5 μg/mL for 120 sec, resulting in a specific interaction between the his-tag and nickel ions. Another baseline step was performed in the assay buffer for 120 sec, after which the ligand-loaded biosensor tips were dipped into the α-syn preparation solutions at different concentrations in 1X PBS (pH 7.4) with 0.05% Tween-20 (PBS-T) for 300 sec. Finally, dissociation was conducted in the assay buffer for 400 sec. A reference biosensor was loaded with the his-tagged sdAb and run with an assay buffer blank for the association and dissociation steps. The biosensor tips were regenerated by cycling them three times for 5 sec each between 10 mM glycine, pH 2, and assay buffer, followed by re-charging for 1 min with 10 mM NiCl_2_.

Data analysis was performed using Data Analysis 11.0 software with reference subtraction using a 1:1 binding model. In a 1:1 bimolecular interaction, both the association and dissociation phases display a time-resolved signal that is described by a single exponential function. Analyte molecules bind at the same rate to every ligand binding site. The association curve follows a characteristic hyperbolic binding profile, with an exponential increase in signal followed by leveling off to a plateau as the binding reaches equilibrium. The dissociation curve follows a single exponential decay with the signal eventually returning to baseline. The complete fitting solution for a 1:1 binding is provided as follows:

Association phase:

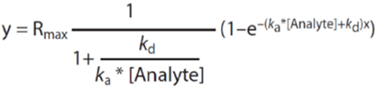

Dissociation phase:

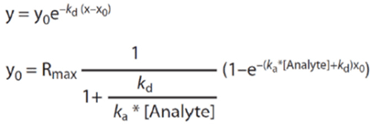

The curve fitting algorithm utilized the Levenberg-Marquardt fitting routine to obtain the best curve fit. In the 1:1 model, the data fits the above equations with x, y, and Analyte representing time, nm shift, and concentration, respectively. The algorithm determines Rmax, k_d_ and k_a_ values, from which K_D_ is calculated by dividing k_d_ by k_a_ (i.e. K_D_ = k_d_/k_a_).

All data were inverted from negative to positive to enable curve fitting of both association and dissociation phases. The ForteBio analytical protocol recommends global analysis of a broad range of analyte concentrations, where the data are analyzed simultaneously by fitting both association and dissociation phases using a 1:1 binding model with the same rate constants. Global analysis enhances the robustness and accuracy of the estimated binding constants. For detailed information, refer to the company’s website (www.sartorius.com) for the Application Note on Biomolecular Binding Kinetics Assays on the Octet® Platform. The Rmax was unlinked by sensor and the data were grouped by color during global fitting.

### Immunoprecipitation (IP) depletion of *α*-syn from LBD brain soluble fraction and dot blot assay

The protein concentration of the LBD brain soluble fraction was determined using a BCA assay and normalized to 1 mg/ml prior to the IP experiments. All procedures were carried out at 4°C to maintain protein stability. The IP was conducted using the Thermo Scientific Pierce Crosslink Magnetic IP/Co-IP Kit (Catalog # 88805) following the manufacturer’s protocol. Initially, the beads were pre-washed with 1X Coupling Buffer. Subsequently, 10 µg of α-syn antibody (211) was diluted to a final volume of 100 µL and immobilized on Protein A/G magnetic beads for 15 minutes. After three washes with 1X Coupling Buffer, the antibody was crosslinked to the beads using disuccinimidyl suberate (DSS) for 30 minutes. Following three washes with Elution Buffer and two washes with IP Lysis/Wash Buffer, the LBD brain soluble fraction (300 µL) containing 300 μg of total protein was incubated with the prepared beads overnight at 4°C. The beads were then separated using a magnetic stand, and the unbound sample, S1 (IP α-syn flow through), was collected for dot blot analysis. After two washes with IP Lysis/Wash Buffer and one wash with purified water, the bound antigen was eluted by incubating the beads with 300 µL of Elution Buffer for 5 minutes at room temperature on a rotator. The beads were magnetically separated, and the supernatant, S1 (IP α-syn), containing the α-syn antigen was retained for dot blot analysis.

For the dot blot assay, 5 µL of S1 (IP α-syn) and S1 (IP α-syn flow through), 5 µL of LBD brain soluble fraction supernatant (S1, 1 mg/ml), rec α-syn (1 mg/ml), and BSA (bovine serum albumin, 1 mg/ml) were spotted onto a nitrocellulose membrane (0.2 μm) and air-dried for 30 minutes. Subsequently, the membrane was blocked for 1 hour with 5% milk in 0.1% Tween-20 in Tris-buffered saline (TBS-T). The blots were then incubated overnight at 4°C with either 2D8 or modified 2D8 (2D8-PEGn-T, n=2,4,6) at a concentration of 0.01 mg/ml in Superblock (Thermo Fisher Scientific). All sdAbs are equipped with a His tag. After several washes in TBS-T, the blots were incubated for 1 hour at room temperature with anti-6×His tag antibody (1:2000, ABCAM, EPR20547) in Superblock, followed by additional washes with TBS-T. The signals were detected using an IRDye 800CW secondary antibody (1:10,000, LI-COR Biosciences). Immunoreactive signals were visualized and quantified using LI-COR Image Studio Lite 5.2.

### Human brain tissue

Unfixed cortical brain samples from a subject with extensive cortical α-syn inclusions indicative of Lewy body dementia were obtained from the National Disease Research Interchange (Philadelphia, PA). These samples were homogenized in Tris buffered saline (TBS) and were centrifuged at 3000 x g for 20 min. The supernatant was retained as the total soluble fraction (S1), while the pellet was resuspended in Tris-EDTA (TE) buffer and labeled as the total insoluble fraction containing the larger α-syn aggregates (P1). The S1 fraction was further centrifuged at 20,000 x g for 20 min. That supernatant was retained as soluble fraction S2, while the pellet was resuspended in TE buffer and labeled as the P2 insoluble fraction. The S2 fraction was further centrifuged at 100,000 x g for 60 min. That supernatant was retained as soluble fraction S3, and the pellet was resuspended in TE buffer and labeled as the P3 insoluble fraction. See **Supplemental Figure 2A** for schematic protocol illustrating the preparation of these different brain homogenate fractions.

Toxicity of these different fractions was initially assessed in primary neuronal cultures that were prepared from wild-type pups at day 0 (**Supplemental Figure 2B-C**). The cultures were incubated for 7 days with 10 μg/ml of each of the six brain fractions. Toxicity was assessed by western blotting for GAPDH. Subsequent experiments used the S1 fraction in primary neuronal cultures that were prepared from transgenic M83 A53T α-syn mice.

### Preparation of primary neuronal cultures

Primary neuronal cultures were prepared from M83 mouse pups at postnatal day 0 according to established protocols (*39*). The homozygous M83 mice express the A53T mutation of human α-syn via the mouse prion promoter at approximately twelve times the level of endogenous mouse α-syn. This model develops extensive α-syn inclusions with age, particularly in the homozygous mice at the age used here (5-8 months), and various behavioral deficits have been reported in this model (*51*). In brief, plates were coated with Pluripro (Cell Guidance Systems) for at least 3 h. The cortex and hippocampus of each mouse were collected and washed in modified Hank’s Balanced Salt Solution. Tissue was then incubated with 0.5% trypsin for 20 min at 37°C, followed by further washing and manual dissociation. Cells were maintained in plating media, whereas the plating media was replaced with neurobasal media to eliminate glia and produce neuronal cultures after the first 24 h.

### *α*-Syn seeding experiment

To determine the seeding capacity, induced by soluble S1 fraction from the human LBD brain, we performed a dot-blot assay using two conformational antibodies, OC and A11, that have been described to bind to various oligomers of different primary amino acid sequences (*52, 53*). Briefly, primary neuronal cultures were prepared from M83 mouse pups as mentioned above. The cultures were incubated for 1, 2, 3, 5, and 7 days with or without 10 μg/ml of soluble S1 fraction. Cell lysate was normalized to 0.5 mg/ml. For the dot blot assay, 5 µL of cell lysates were spotted onto a nitrocellulose membrane (0.2 μm) and air-dried for 30 minutes. Subsequently, the membrane was blocked for 1 hour with 5% milk in 0.1% Tween-20 in Tris-buffered saline (TBS-T). The blots were then incubated overnight at 4°C with conformational antibodies OC (1:1000, Millipore #AB2286) or A11 (1:1000, Thermo Fisher #AHB0052) in Superblock (Thermo Fisher Scientific). The signals were detected using an IRDye 800CW secondary antibody (1:10,000, LI-COR Biosciences). Immunoreactive dots were visualized and quantified using LI-COR Image Studio Lite 5.2.

### Cell Culture Efficacy and Degradation Mechanisms

To evaluate the efficacy of each sdAb in clearing pathological α-syn or preventing α-syn-induced toxicity and seeding, we tested them in primary neuronal cultures in two dosing paradigms that simulate extracellular and intracellular mechanisms.

For the extracellular blockage of α-syn spreading, we added LBD brain-derived α-syn (S1 fraction, 10 μg/ml) and sdAbs (5 μg/ml of 2D8 or molar equivalent of modified 2D8 - about 3% weight difference) to the culture simultaneously (S1 + sdAb) and incubated them together in the media for 24 h. The media was replaced, and the cells were maintained in culture for an additional 96 h. Samples were collected 96 h after antibody application.

Intracellular clearance was modeled by adding LBD brain-derived α-syn (S1 fraction, 10 μg/ml) to the neuronal cultures first and allowing the cells to take up the pathological protein for 24 h. We then exchanged the media and added different concentrations of sdAbs ranging from 0.625-10 µg/ml of 2D8 or molar equivalent of modified 2D8 (S1 → sdAb). Since the soluble S1 α-syn fraction was already internalized in this paradigm, the sdAb must also enter the cells to be effective. Samples were collected 96 h after antibody application.

For all experiments, we collected a set of Day 0 untreated controls from the same plate before treatment. Additionally, an untreated group of cells was collected together with the experimental groups once the incubation was complete. All groups were compared to their respective Day 0 untreated controls to ensure that changes seen in the tests were not the result of spending an additional 4 days in culture. Moreover, all samples on the same plate were prepared from the same animal to control for any variation in protein expression between mice and the overall age of the culture.

To determine the degradation mechanism of sdAbs, we incubated M83 primary neurons with 10 µg/ml of LBD brain-derived soluble S1 α-syn fraction for 24 h. We then exchanged the media, pre-treated the neurons with either lysosome inhibitor Baf.A1 (0.5 or 1 µM) or proteasome inhibitor MG132 (5 or 10 µM) for 6 h, and added 5 µg/ml unmodified sdAb 2D8 or modified sdAb 2D8-PEG4-T directly into the media without exchanging it. Samples were collected two days after antibody application.

### Immunofluorescence assay

For imaging, transgenic M83 primary neuronal cultures were prepared on glass bottom plates as previously described, and we examined the uptake of sdAb and its co-localization with intracellular α-syn. Neurons were first treated with 10 µg/ml LBD brain-derived soluble S1 α-syn fraction for 24 h, and then the media was exchanged, and 5 µg/ml of either unmodified sdAb 2D8 or modified sdAb 2D8-PEG4-T were added and incubated for another 2 h. After that, the neuron culture was washed with PBS, fixed with 4% (v/v) formaldehyde-HBSS containing 4% sucrose for 20 min at room temperature, washed with HBSS for 10 min three times, and then incubated with primary antibodies (HIS.H8 and total α-syn 211) in HBSS containing 5% BSA and 1% normal donkey serum solution at 4°C overnight. After washing with HBSS, the bound antibodies were detected with an Alexa Fluor 488 goat anti-rabbit IgG or Alexa Fluor 568 goat anti-mouse IgG (Invitrogen), and the nuclei were labeled with DAPI in HBSS containing 5% BSA and 1% normal donkey serum solution for 1 h at room temperature. Finally, the samples were covered with ProLong Gold, and images were obtained using an LSM 800 Zeiss confocal laser scanning microscope (Axioimager).

To measure the co-localization of sdAb and α-syn, the Manders’ correlation coefficient was calculated using the JACop plugin in Image J software (*54*). The Manders’ coefficients represent the proportion of confocal fluorescence signals, where 0 is defined as no co-localization and 1 is defined as complete co-localization.

### Co-immunoprecipitation assays

Primary neuron cultures intended for co-immunoprecipitation (co-IP) assays were prepared following the procedures described above. M83 primary neurons were then exposed to 10 µg/mL LBD brain-derived soluble S1 α-syn fraction for 24 h to allow the cells to take up the pathological α-syn. Media was then exchanged, and the neurons were pre-treated with 10 µM of proteasome inhibitor MG132 for 6h. Unmodified sdAb 2D8 or modified sdAb 2D8-PEG4-T at 5 µg/mL were directly added into the media without media exchange, and samples were collected 2 days after the antibody application. Cell lysis was performed with ice-cold Pierce IP Lysis Buffer (ThermoFisher Scientific) for 15 min at 4°C, followed by centrifugation at 10,000 g for 10 min. The supernatant (Input) was transferred to a new tube and immediately aliquoted for the BCA assay to determine protein concentration. All steps were conducted at 4°C to stabilize and detect complex formation. Co-IP was performed using the Thermo Scientific Pierce Crosslink Magnetic IP/Co-IP Kit (Catalog # 88805) according to the manufacturer’s instructions. First, the beads were pre-washed with 1X Coupling Buffer. Then, 10 µg α-syn antibody (211) or control IgG antibody were diluted to a final volume of 100 µL and bound to Protein A/G magnetic beads for 15 min. The beads were washed three times with 1X Coupling Buffer, and the antibody was cross-linked to the beads with disuccinimidyl suberate (DSS) for 30 min. The beads were then washed three times with Elution Buffer and twice with IP Lysis/Wash Buffer. The lysate solution was diluted with IP Lysis/Wash Buffer to 200 µL with a total protein concentration of 300 μg and incubated with prepared beads overnight at 4°C. The beads were washed twice with IP Lysis/Wash Buffer and then once with purified water. The bound antigen was eluted with 100 µL of Elution Buffer and incubated for 5 min at room temperature on a rotator. The beads were magnetically separated, and the supernatant containing the target antigen was saved. The supernatant was diluted with 4x Laemmli Sample buffer (Bio-Rad), and 15 μL of sample was loaded onto SDS-PAGE for western blotting.

### Mouse models, injections of sdAb, and IVIS imaging

#### M83 Mouse Model

Homozygous M83 mice were utilized in this study ((*51*) and JAX# 004479). These mice express the A53T mutation of human α-syn via the mouse prion promoter at approximately twelve times the level of endogenous mouse α-syn. The M83 model exhibits age-dependent development of α-syn inclusions, which are particularly extensive in the homozygous mice at the specific age range employed in this study (7-8 months).

#### In Vivo Imaging System (IVIS) Imaging

Prior to and following sdAb administration, imaging of α-syn lesions was performed using the In Vivo Imaging System (IVIS, Lumina XR, Perkin Elmer) in the homozygous M83 α-syn A53T mice. For detailed information on age and sex, refer to **Supplemental Table 2**. Before IVIS imaging, the mice were anesthetized with 2% isoflurane and maintained with 1.5% isoflurane in 30% oxygen. Subsequently, they were injected intravenously (i.v.) into the femoral vein with VivoTag 680XL (Perkin Elmer) labeled sdAb 2D10 at a dosage of 10 mg/kg. The injection procedure followed established protocols described in previous studies involving this and other substances (*37, 55, 56*). To minimize non-specific light diffraction from hairs, the fur on the head and body was shaved off. Immediately after the i.v. sdAb injection, the mice were imaged at defined intervals of 15 minutes. They remained anesthetized within the imaging chamber and were supplied with 2% isoflurane delivered with oxygen via a nosecone at a flow rate of 0.4 L/min. Their body temperature was maintained at 37°C using a thermoelectrically controlled imaging stage. Fluorescent images were acquired using Living Image software (PerkinElmer) with the following settings: mode: fluorescent, illumination: epi-illumination, exposure time and binning: auto (30 s under these conditions), excitation filter: 675 nm, emission filter: Cy5.5 (695–770 nm). Image data were automatically time and date stamped to facilitate analysis.

#### Analysis of IVIS Images

The acquired images were analyzed using Living Image software from PerkinElmer. First, images from different time points for each mouse were loaded together as a sequence. Subsequently, the images were adjusted to the same color scale to facilitate visual comparison, and a circular region of interest (ROI) was drawn in the brain region of each animal. The calibrated unit of radiant efficiency (photons/sec/cm^2^/Str/mW/cm^2^) was recorded for each region of interest (ROI), and a table containing all ROI values in the sequence was generated, saved, and utilized for graphing the IVIS imaging profile of each mouse.

#### Intravenous Injections

Intravenous injections of PBS, sdAb 2D8, and modified sdAb 2D8-PEG4-T were performed three times into the femoral vein of the M83 mice on Day 4, 7, and 10 (**Figure 7A**). Each injection involved a dosage of 3 mg/kg of the unmodified sdAb and its molar equivalent of the modified sdAb (about 3% weight difference).

**Figure 1:**
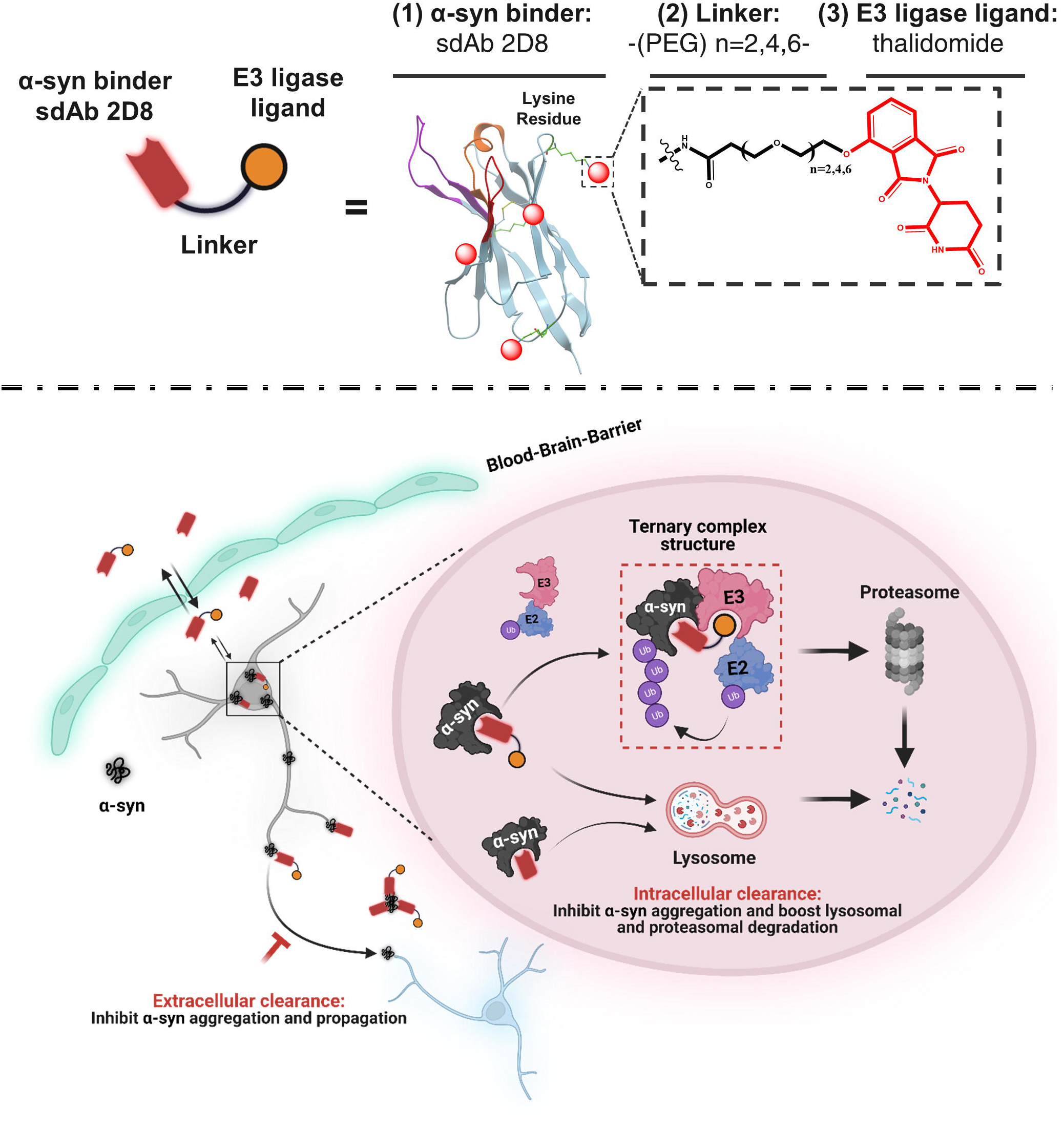
Design and working model for a single-domain antibody (sdAb)-based protein degrader targeting synucleinopathies. The degrader molecule is composed of anti-α-syn sdAb 2D8 connected via a variable-size linker to thalidomide, an E3-ligase recruiting ligand that engages with CRL4^CRBN^. This degrader molecule can enter the mouse brain after i.v. injection and target α-syn both extracellularly and intracellularly. Specifically, extracellularly, the sdAb may sequester α-syn aggregates and disrupt their assembly, and thereby collectively prevent the spread of α-syn pathology between neurons. Intracellularly, the protein degrader may bind to α-syn aggregates within the endosomal-lysosomal system and facilitate their disassembly, which would result in better access of lysosomal enzymes to degrade the aggregates. The degrader molecule also brings α-syn and E3 ligase into proximity to form a ternary complex, which is predicted to mediate α-syn ubiquitination and trigger its degradation by the proteasome. This working model provides a framework for the development of protein degraders for the treatment of synucleinopathies. Abbreviations: α-syn = α-synuclein, E3 = E3 ligase, E2 = E2 ligase, Ub = ubiquitin.

**Figure 2:**
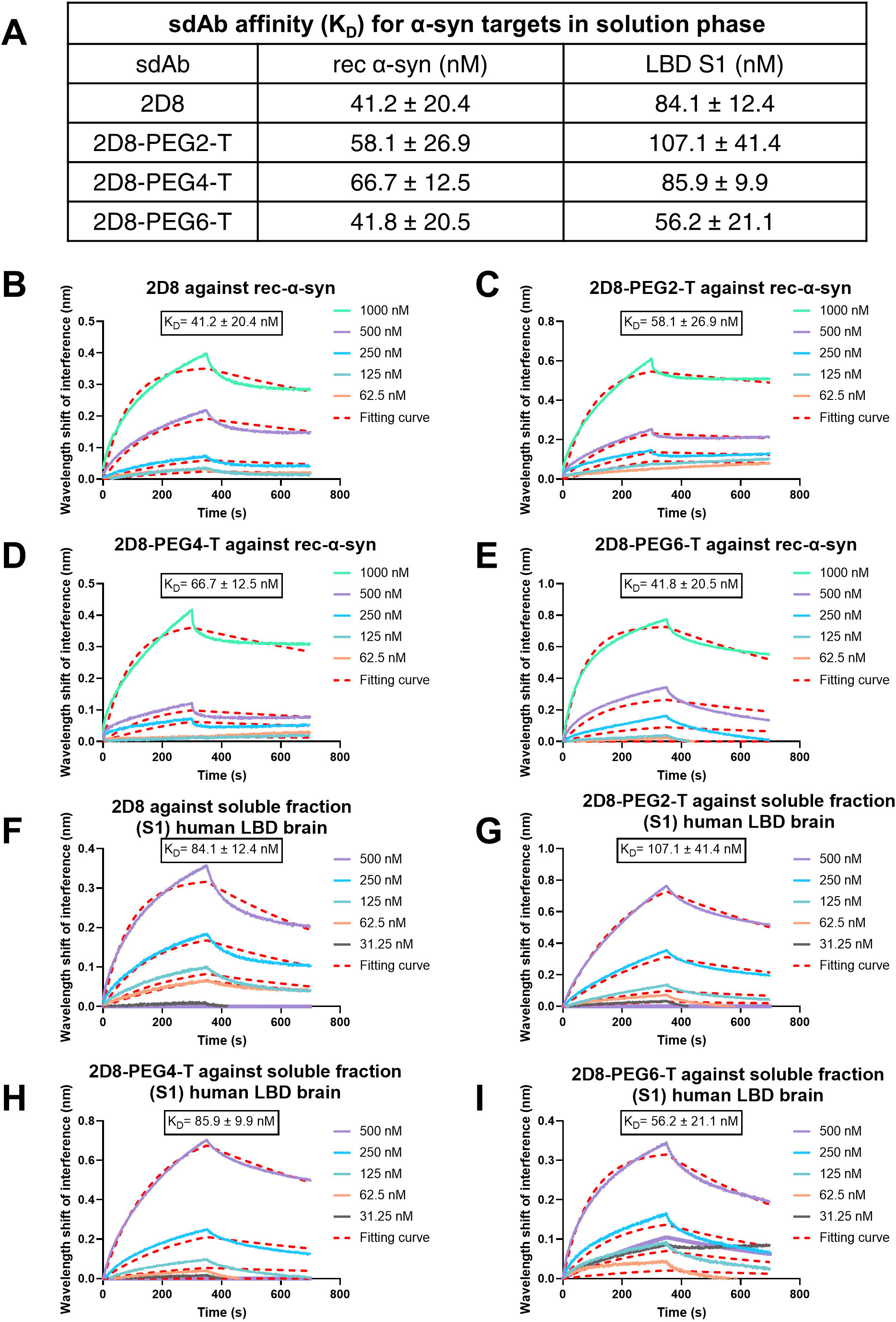
Affinities of unmodified sdAb 2D8 and modified sdAb 2D8-PEGn-T for different *α*-syn preparations. (A) Summary table of binding affinities: To conduct biolayer interferometry assay in solution phase, unmodified sdAb 2D8 and modified sdAbs 2D8-PEGn-T (n=2, 4, 6) were loaded onto the Ni-NTA biosensor, and K_D_ values were determined using increasing concentration of the different α-syn preparations, including rec α-syn and soluble fraction (S1) from LBD brain. (B-I) The representative curves illustrate the wavelength shift of interference in nanometers (nm), which is interpreted as binding. The curves demonstrate the association and dissociation of sdAb and the α-syn preparations at various concentrations, with the broken line indicating the fitting curve used to calculate the K_D_ value ± standard deviation (SD). The K_D_ value was obtained from three independent experiments. Additional information on association (k_a_) and dissociation (k_d_) values can be found in **Supplemental Table 1.**

**Figure 3:**
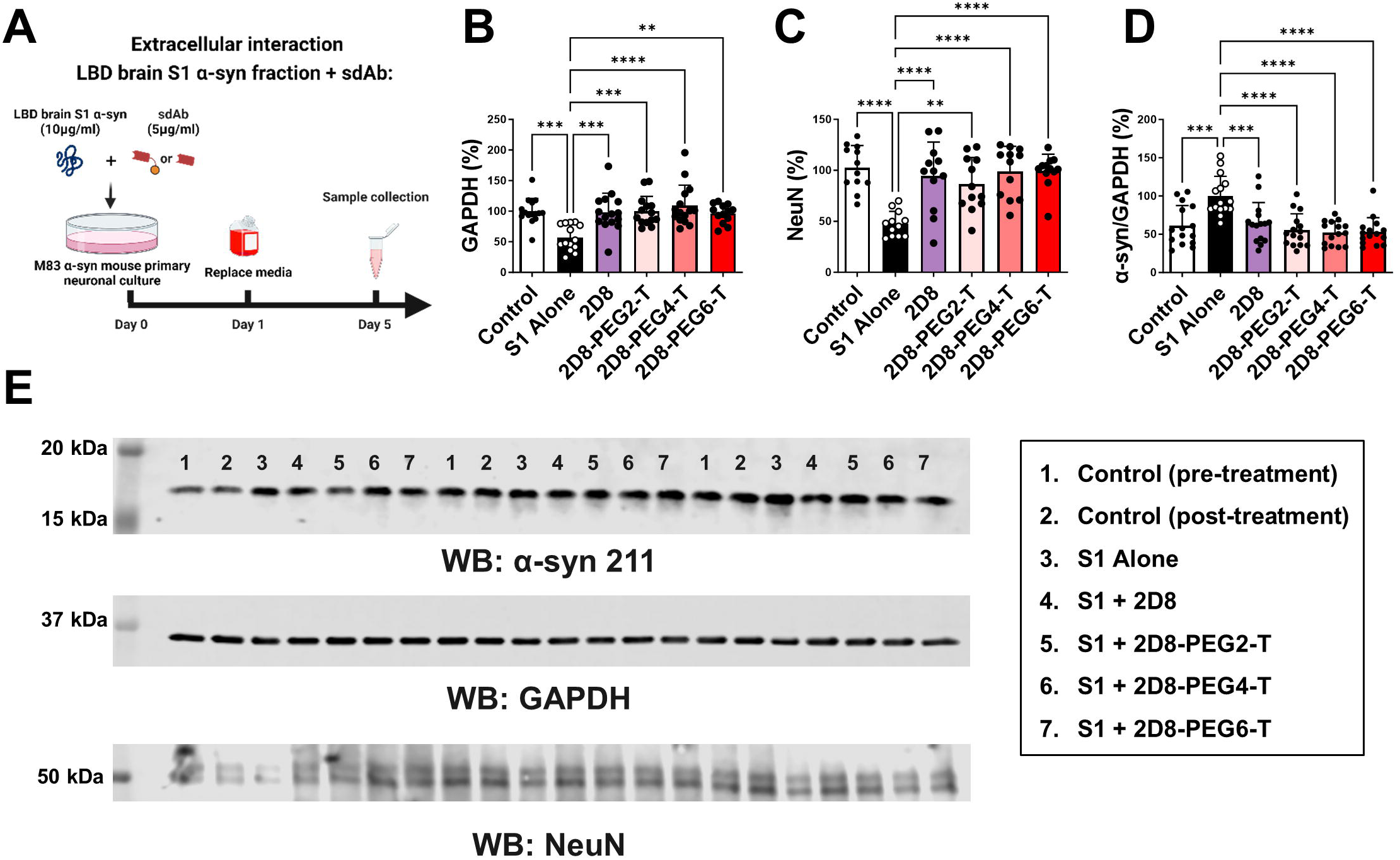
Neuroprotective effects of unmodified sdAb 2D8 and modified sdAb 2D8-PEGn-T. (A) M83 primary neurons were incubated with 10 µg/ml LBD S1 fraction and 5 μg/ml of either 2D8 or its molar equivalent of 2D8-PEGn-T (n=2, 4, 6) simultaneously for 24h. After media replacement, cells were cultured for an additional 96h, with samples collected for analysis (n=at least 12 replicates per condition). (B) One-way ANOVA analysis revealed significant group differences in GAPDH levels (p<0.0001). S1 alone decreased GAPDH levels by 43% compared to control at 96h (p=0.0003). Both modified 2D8 and modified 2D8-PEGn-T (n=2, 4, 6) prevented S1-induced toxicity, increasing GAPDH levels by 42% (p=0.0005, 2D8), 43% (p=0.0003, 2D8-PEG2-T), 52% (p<0.0001, 2D8-PEG4-T), and 39% (p=0.0018, 2D8-PEG6-T) relative to S1 alone, respectively. (C) One-way ANOVA analysis revealed significant group differences in NeuN levels (p<0.0001). S1 alone decreased NeuN levels by 54% compared to control at 96 h (p<0.0001). Both unmodified 2D8 and modified 2D8-PEGn-T (n=2, 4, 6) prevented S1-induced toxicity, resulting in increased GAPDH levels by 50% (p<0.0001, 2D8), 45% (p=0.0017, 2D8-PEG2-T), 52% (p<0.0001, 2D8-PEG4-T), and 52% (p<0.0001, 2D8-PEG6-T) relative to S1 alone, respectively. (D) One-way ANOVA analysis revealed significant group differences in total α-syn/GAPDH levels (p<0.0001). S1 alone increased α-syn levels by 39% compared to control at 96 h (p=0.0002). Both sdAb 2D8 and modified sdAbs 2D8-PEGn-T (n=2, 4, 6) prevented α-syn seeding, resulting in decreased total α-syn/GAPDH levels by 35% (p=0.0008, 2D8), 45% (p<0.0001, 2D8-PEG2-T), 48% (p<0.0001, 2D8-PEG4-T), and 47% (p<0.0001, 2D8-PEG6-T) relative to S1 alone, respectively. (E) Immunoblots illustrating total α-syn, GAPDH, and NeuN levels in treated M83 cell lysate. The complete western blots and bands analyzed are shown in **Supplemental Figure 5.** **, ***, ****: p<0.01, 0.001, 0.0001 (Tukey’s post-hoc test).

**Figure 4:**
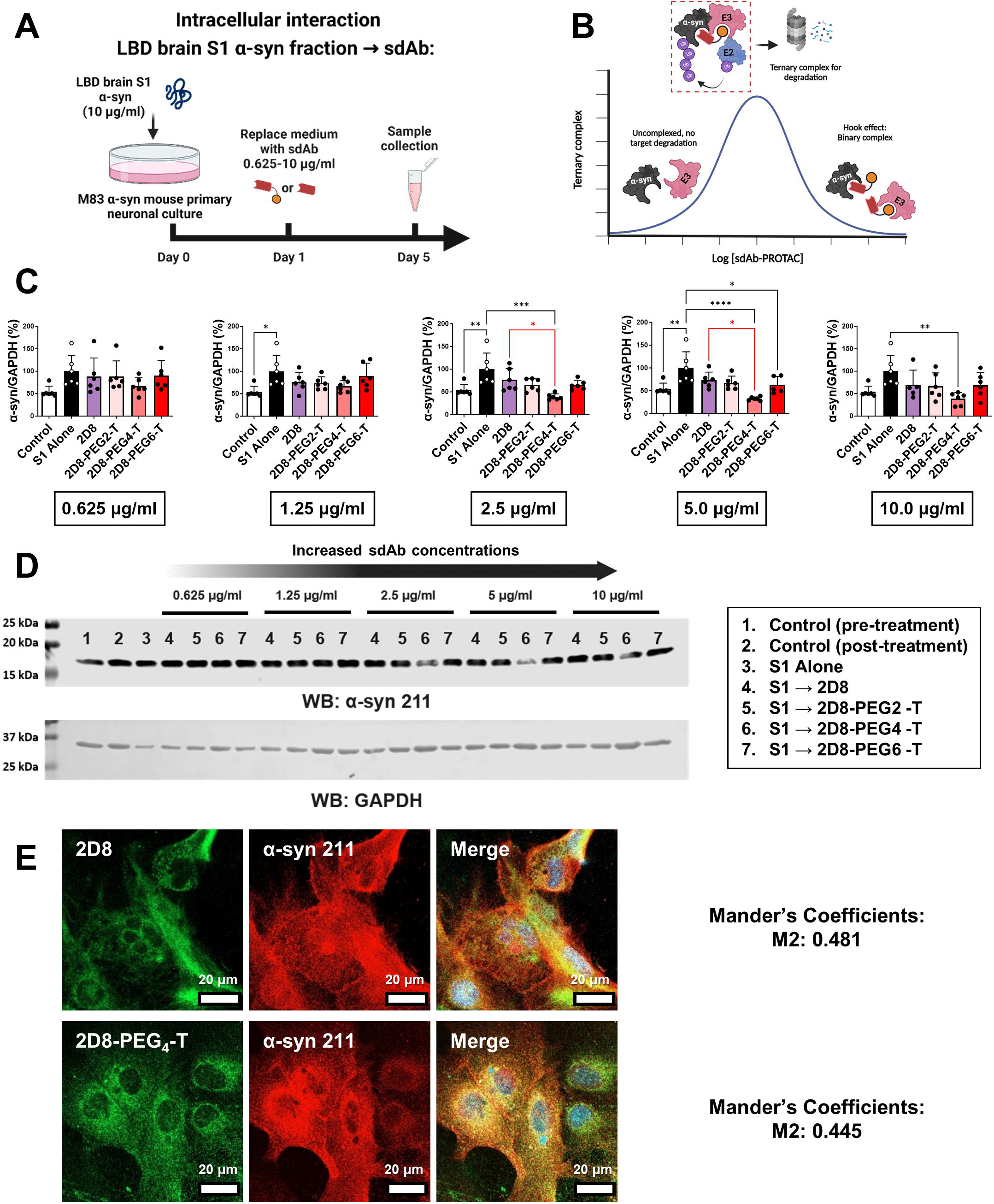
Enhanced intracellular clearance of *α*-syn by modified sdAb 2D8-PEG4-T in primary neurons. (A) M83 primary neurons were incubated with 10 µg/ml LBD S1 fraction for 24h. Subsequently, different concentrations of sdAbs ranging from 0.625-10 µg/ml of 2D8 and molar equivalent of modified 2D8 were added, with samples collected 96h post-antibody application. (B) Schematic diagram of PROTAC-mediated ternary complex formation and Hook effect based on PROTAC concentration. (C) One-way ANOVA analysis revealed significant differences in normalized total α-syn levels when sdAb concentration exceeded 0.625 µg/ml (p=0.02 for 1.25 µg/ml, p=0.0004 for 2.5 µg/ml, p=0.0001 for 5.0 µg/ml, p=0.0136 for 10.0 µg/ml). At 1.25-5.0 µg/ml sdAb concentration, S1 alone increased α-syn levels by 46% compared to control at 96h (p=0.0154 (1.25 µg/ml), p=0.0049 (2.5 µg/ml), p=0.0043 (5.0 µg/ml)). At 2.5-5.0 µg/ml, 2D8-PEG4-T reduced S1-induced increase in total α-syn levels (reduced by 61% at 2.5 µg/ml, p=0.0001, and 67% at 5.0 µg/ml, p<0.0001, relative to S1 alone). Modified sdAb 2D8-PEG4-T also lowered total α-syn levels compared to unmodified 2D8 at 2.5 and 5.0 µg/ml (reduced by 38% at 2.5 µg/ml, p=0.0298, and 41% at 5.0 µg/ml, p=0.0126, relative to unmodified 2D8). (D) Immunoblots depicting total α-syn and GAPDH levels in treated M83 cell lysate. Complete western blots and bands analyzed are in **Supplemental Figure 6.** (E) Immunofluorescence of M83 primary mouse neurons treated with 10 µg/ml LBD S1 fraction for 24h. Media was then exchanged, and unmodified sdAb 2D8 or modified sdAb 2D8-PEG4-T at 5.0 µg/ml or molar equivalent were added and incubated for an additional 2h. Neurons were subsequently immuno-probed for sdAb containing His tag (HIS.H8, green) and total α-syn (211, red). Scale bar: 20 μm. Mander’s coefficient M2 quantifies the ratio of α-syn overlapping with sdAb. Calculated as the sum of overlapping intensity in α-syn divided by total intensity of α-syn. *, **, ***, ****: p<0.05, 0.01, 0.001, 0.0001 (Tukey’s post-hoc test).

**Figure 5:**
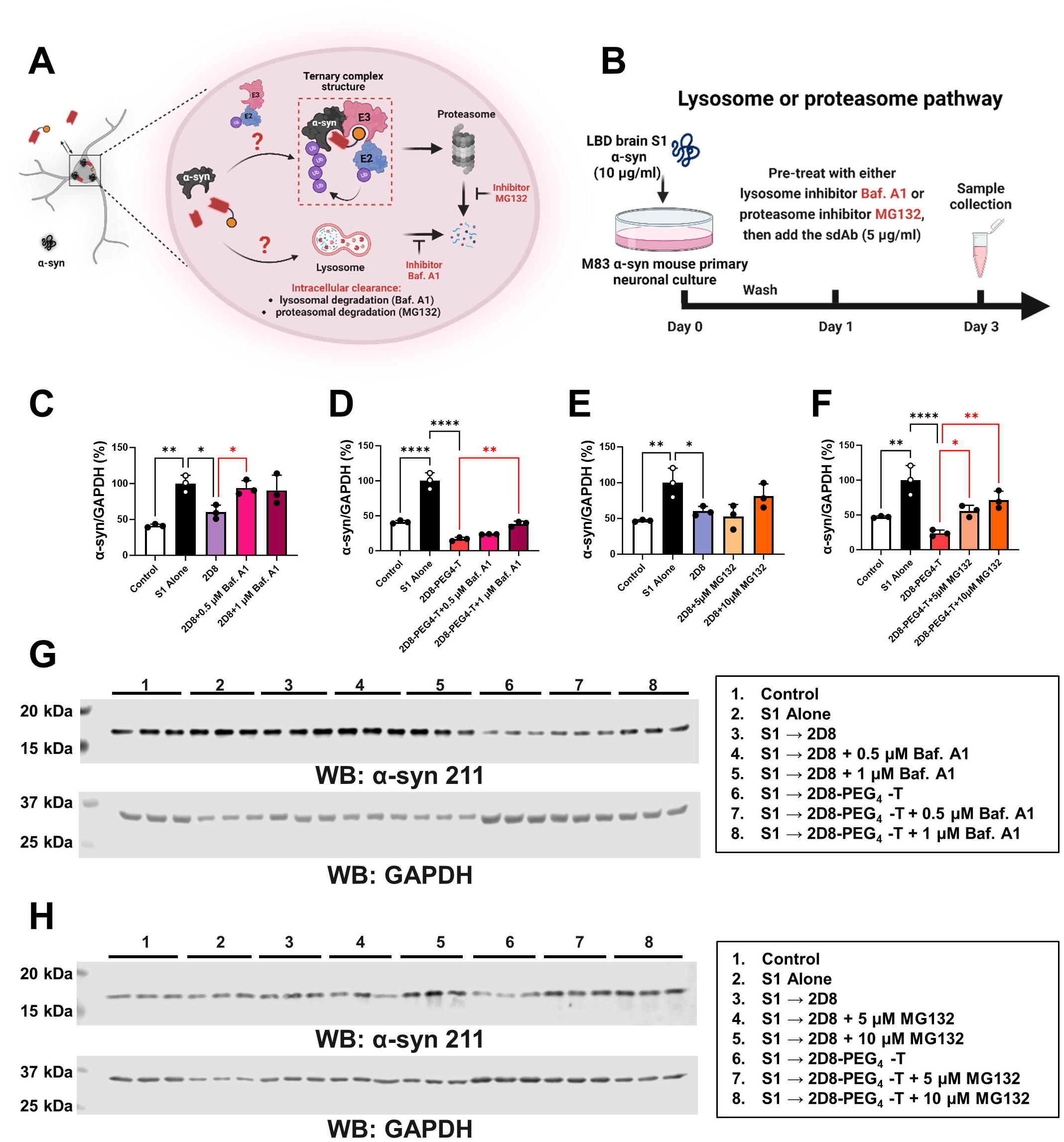
*α*-Syn clearance by unmodified and modified sdAbs via lysosome and proteasome pathways. (A) To determine the degradation mechanism of 2D8 and 2D8-PEG4-T, the lysosome and proteasome pathways were blocked using bafilomycin A1 (Baf.A1) and MG132 inhibitors, respectively. (B) M83 primary neurons were incubated with 10 µg/ml LBD S1 fraction for 24h to allow uptake of pathological α-syn. The media was then replaced, and the neurons were pre-treated with either the Baf.A1 or MG132 for 6h. Subsequently, the neurons were treated with 5 µg/ml 2D8 or molar equivalent of modified 2D8 for 2-days. (C-F) One-way ANOVA analysis revealed group differences in normalized total α-syn levels (p=0.0007 (C), p<0.0001 (D), p=0.0060 (E), and p=0.0002 (F)), N=3. For the Baf.A1 inhibitor (C-D), S1 alone increased α-syn levels by 59% compared to control (p=0.0013 (C), p<0.0001 (D)). Both 2D8 and 2D8-PEG4-T prevented S1-induced increase in total α-syn levels (reduced by 40% for 2D8, p = 0.0194 (C); and 83% for 2D8-PEG4-T, p<0.0001 (D)). However, the Baf.A1 inhibitor blocked this degradation effect, leading to a 34% increase for 2D8 (p=0.0494) with 0.5 µM of inhibitor and a 22% increase for 2D8-PEG4-T (p=0.0054) with 1 µM of inhibitor (C-D). For the MG132 inhibitor (E-F), S1 alone increased α-syn levels by 53% compared to control (p=0.0077 (E), p=0.0018 for (F)). Both 2D8 and 2D8-PEG4-T prevented the S1-induced increase in total α-syn levels (reduced by 40% for 2D8, p = 0.0434 (E), and 76% for 2D8-PEG4-T, p<0.0001 (F)). The MG132 inhibitor did not block α-syn clearance of 2D8 (E), but it blocked α-syn clearance of 2D8-PEG4-T in a concentration-dependent manner (F), resulting in a 32% increase and 48% increase in α-syn with 5 µM (p=0.0437) and 10 µM MG132 (p=0.0040) for 2D8-PEG4-T. (G-H) Immunoblots show total α-syn and GAPDH levels for C-F. Complete western blot analysis are in **Supplemental Figure 7.** *, **, ****: p<0.05, 0.01, 0.0001 (Tukey’s post-hoc test).

**Figure 6:**
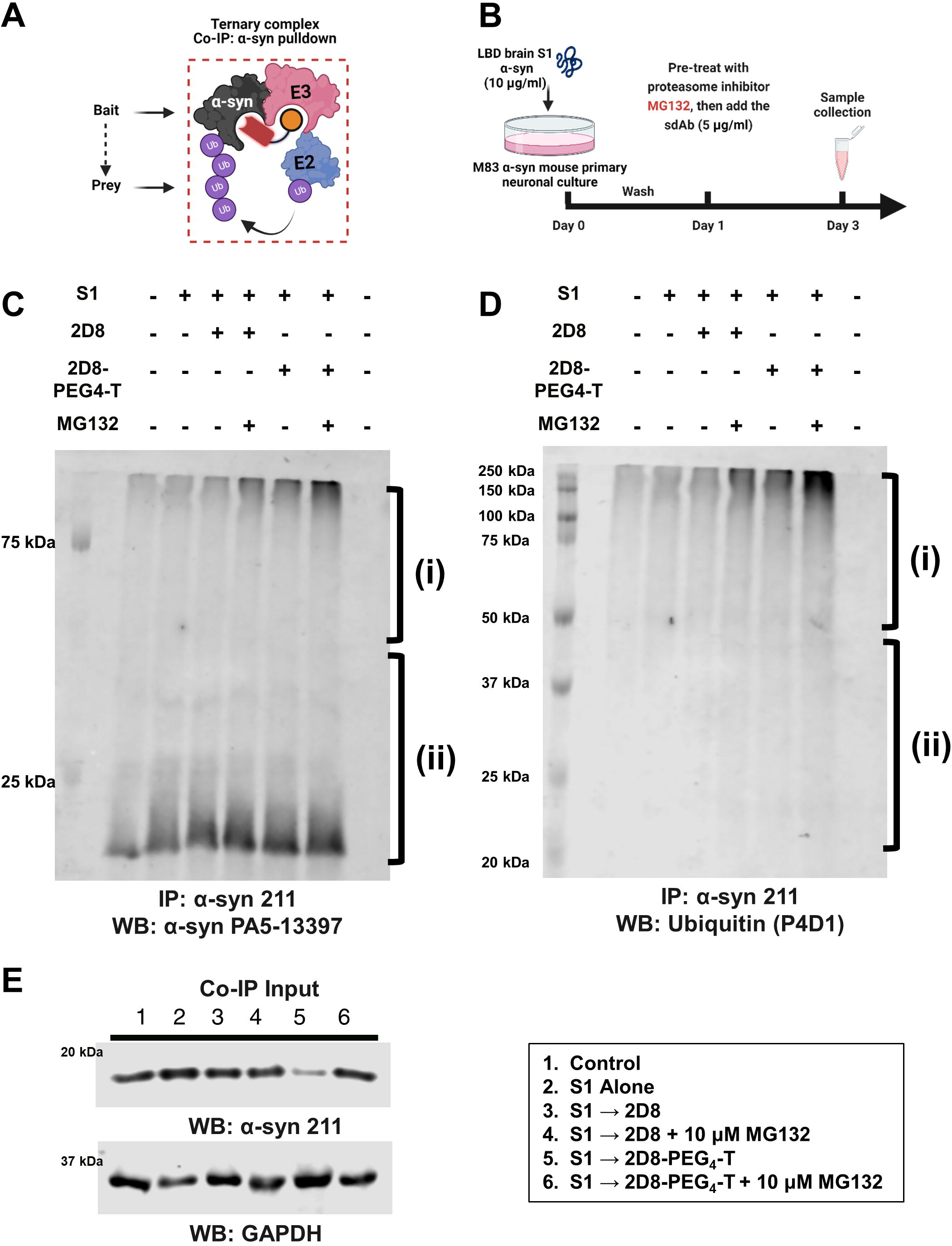
Ternary complex formation and sdAb-mediated effects on pathological *α*-syn in M83 primary neurons. (A) Co-immunoprecipitation (co-IP) and western blot analysis demonstrate the formation of ternary complexes in M83 primary neurons upon sdAb treatment. (B) Experimental procedure: M83 primary neurons were treated with 10 µg/ml LBD brain soluble fraction (S1) for 24 h to allow uptake of pathological α-syn. The media was then exchanged, and the neurons were pre-treated with the proteasome inhibitor MG132 (10 µM) for 6 h. Next, sdAb 2D8 or sdAb 2D8-PEG4-T (5 µg/ml or its molar equivalent) were added directly into the media without media exchange. Samples were collected 2 days after antibody application. (C) Co-IP using α-syn pulldown (α-syn 211 antibody) was performed, followed by detection of α-syn (PA5-13397 antibody). Long (i) and short (ii) conjugation of the antibody with α-syn is shown. (D) Co-IP using α-syn pulldown (α-syn 211 antibody) was performed, followed by the detection of ubiquitinated proteins (P4D1 antibody). Long (i) and short (ii) ubiquitinated chain conjugation with α-syn is shown. (E) Control western blot analysis with IP input confirms the effect of modified sdAb 2D8-PEG4-T on α-syn with or without the proteasome inhibitor MG132. Complete western blot analysis can be found in **Supplemental Figure 8.**

**Figure 7:**
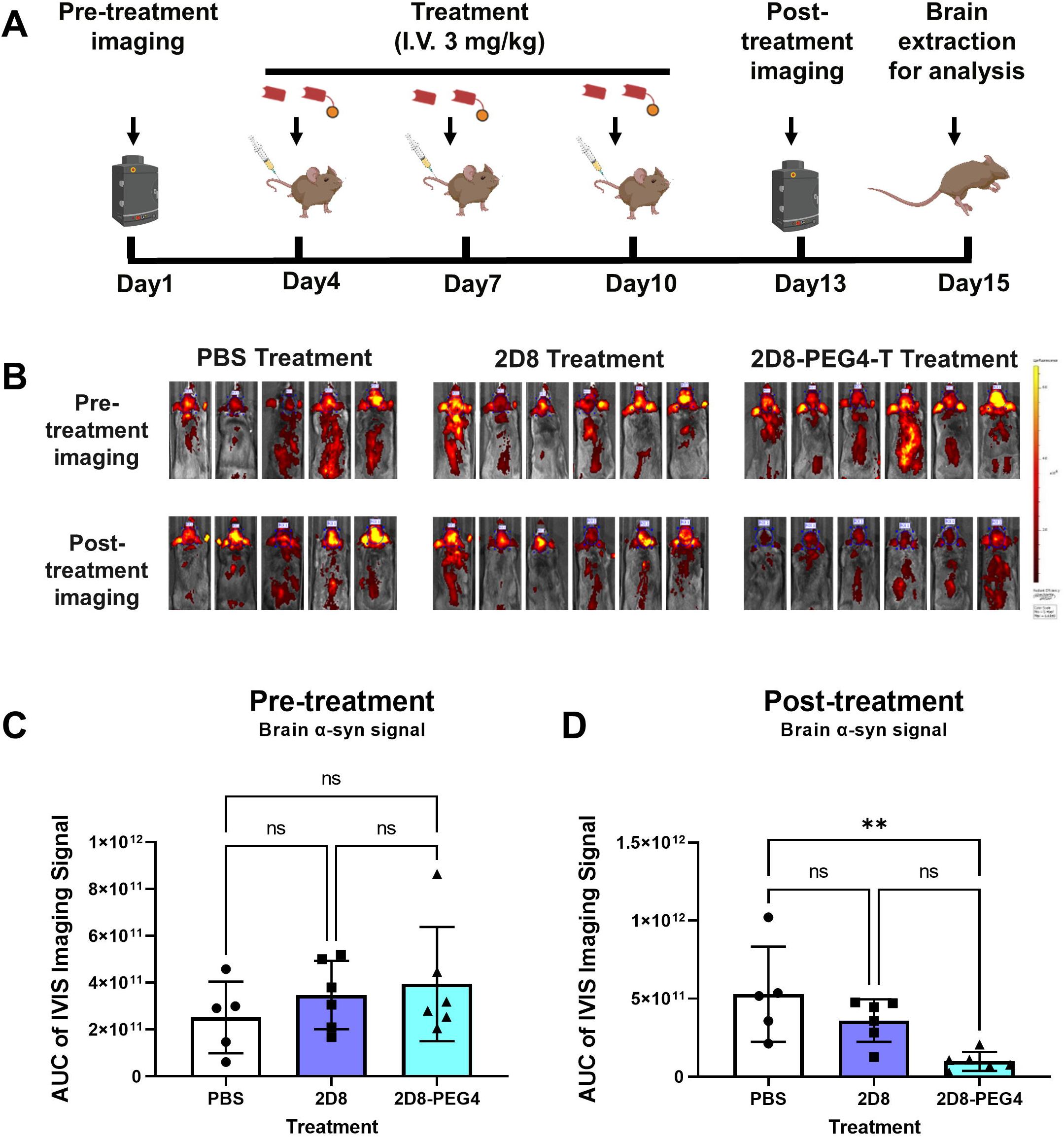
*α*-syn brain signal following anti-*α*-syn sdAb administration in M83 synucleinopathy mice. (A) Schematic diagram illustrating the protocol for anti-α-syn sdAb administration in M83 synucleinopathy mice. To evaluate initial α-syn burden, all mice received an i.v. injection of fluorescently labeled anti-α-syn sdAb 2D10 (10 mg/kg), and in vivo imaging was performed using the IVIS system. Subsequently, the mice received i.v. injections of either PBS, sdAb 2D8 (3 mg/kg), or sdAb 2D8-PEG4-T (molar equivalent of 3 mg/kg 2D8) on days 4, 7, and 10. On day 13, the mice were injected with fluorescently labeled 2D10 again (10 mg/kg) to reassess α-syn burden. The mice were then perfused, and their brains extracted for tissue analyses. (B) Representative images depicting the brain signal intensity after i.v. injection of labeled sdAb-2D10 (10 mg/kg) before and after sdAb administration. The scale bar represents the maximum pixel intensity. Quantitative analysis of the brain signal was conducted by calculating the total radiant efficiency of the summed pixel intensity, derived from the area under the curve (AUC) cumulatively within 2 h post-injection (C-D). (C) Mice were assigned to groups with similar brain α-syn signal as confirmed by one-way ANOVA analysis indicating non-significant (ns) group differences in cumulative brain α-syn signal before sdAb administration (p=0.4676). (D) One-way ANOVA revealed group differences in cumulative brain α-syn signal after sdAb administration (p=0.0059). Compared to the PBS group, sdAb 2D8-PEG4-T decreased brain α-syn signal by 81% (**: p=0.0049).

### Brain homogenization, preparation of *α*-syn fractions and Western blots

#### Brain Collection and Homogenization

After imaging, the brains were collected for biochemical and immunohistochemistry assays. The left hemisphere of the mouse brains was homogenized in modified RIPA buffer (50 mM Tris-HCl, 150 mM NaCl, 1 mM EDTA, 1% Nonidet P-40, pH 7.4) with the addition of protease and phosphatase inhibitors (1X protease inhibitor mixture (cOmplete, Roche), 1 mM NaF, 1 mM Na_3_VO_4_, 1 nM PMSF, and 0.25% sodium deoxycholate). The homogenate was briefly kept on ice and then centrifuged at 20,000 x g for 20 minutes at 20°C. The resulting supernatant was collected as the soluble α-syn fraction (Low-Speed Supernatant, LSS). The protein concentration of each sample was determined using the BCA assay, and the samples were adjusted to the same concentration with the homogenization buffer. Subsequently, 5 μg of protein was loaded per well.

#### Preparation of Sarkosyl Insoluble Fraction

To obtain the sarkosyl insoluble fraction, an equal amount of protein (2 mg) from the LSS was mixed with 10% sarkosyl in phosphate buffer to achieve a final concentration of 1% sarkosyl. The mixture was incubated for 30 min at room temperature on a Mini LabRoller (Labnet H5500), followed by ultracentrifugation at 100,000 x g for 1 h at 20 °C using a Beckman Coulter OptimaTM MAX-XP Ultracentrifuge. The resulting pellet was washed with a 1% sarkosyl solution and centrifuged again at 100,000 x g for 1 h at 20 °C. The sarkosyl pellet (SP) fraction was air dried for 30 min and then mixed with 50 μL of modified O+ buffer (62.5 mM Tris-HCl, 10% glycerol, 5% β-mercaptoethanol, 2.3% SDS, 1 mM EDTA, 1 mM EGTA, 1 mM NaF, 1 mM Na_3_VO_4_, 1 nM PMSF, and 1X protease inhibitor mixture). Similarly, the LSS was eluted with O+ buffer (1:5). All samples were boiled, and 10 μL of each sample was electrophoresed on 12% SDS-PAGE gels.

#### Western Blot Analysis

Following protein transfer, all blots for α-syn detection were treated for 30 min with 4% paraformaldehyde to prevent α-syn from washing off the membrane (*57*). The blots were then blocked in Superblock® (Thermo Fisher Scientific) and incubated with the following primary antibodies: α-syn 211 antibody (1:1000, Invitrogen), Phospho-α-syn (Ser129) antibody (1:1000, D1R1R clone, Cell Signaling Technology), ubiquitin (P4D1, Cell Signaling Technology) (1:1000), and GAPDH (1:4000, Cell Signaling Technology, D16H11). Subsequently, the blots were incubated with either IRDye® 800CW or IRDye® 680RD secondary antibodies (1:10,000, LI-COR Biosciences). Immunoreactive bands were visualized and quantified using LI-COR Image Studio Lite 5.2.

### Immunocytochemistry

The right-brain hemisphere was cut coronally into 40 µm thick sections, using a freezing cryostat, and stored at −20°C in cryoprotectant (20% glycerol in 2% DMSO in 0.1 M phosphate buffer, pH 7.4) until retrieved for immunohistochemistry. After several washes in PBS to remove the cryoprotectant, tissue sections were placed in a free-floating staining rack with 0.3% hydrogen peroxide in Tris buffered saline (TBS, pH 7.4) with 0.25% Tween 20 to block endogenous peroxidases and to permeabilize the sections. Non-specific binding sites were then blocked for 1 h at room temperature using 5% powdered milk in TBS. The primary incubation was performed overnight at 4°C (phospho-Ser129, BioLegend Cat # 825701, 1:2000; GFAP, DAKO Cat # GA524, 1:1000; Iba-1, WAKO Cat # 019-19741, 1:1000). The following day, tissues were washed with TBS containing 0.05% Tween 20. The secondary antibody (1:10,000 M.O.M. Biotinylated Anti Mouse Cat # MKB-2225-.1 or 1:1000 Goat Anti Rabbit IgG, Vector Laboratories Cat # BA-1000) was incubated for 2 h with 20% Superblock (Fisher Scientific Cat # 37535). To enhance the signal intensity, tissues were incubated in an avidin-peroxidase solution (Vectastain Elite Cat # PK-6200). Subsequently, tissues were washed twice with 0.2 M sodium acetate, pH 6.0 for 10 min. Tissues were then transferred to the chromogen solution (0.3% hydrogen peroxide, 0.05% 3, 3 ′-diaminobenzidine tetrahydrochloride, and 0.05% nickel ammonium sulfate hexahydrate in 0.2 M sodium acetate, pH 6). After achieving color development, tissues were washed twice with 0.2 M sodium acetate for 10 min and mounted from a PBS solution on positively charged microscope slides (Fisher Scientific Cat # 15-188-48). Once mounted and dried, slides underwent dehydration through Citrisolv and a series of alcohol gradients (70, 80, 95, and 100%) and were coverslipped (Fisher Scientific Cat 12545F) with DEPEX mounting medium (Electron Microscopy Sciences Cat # 13515).

The sections on slides were imaged at 20X using a Leica 5000B microscope, and images were captured with Stereo Investigator (Version 2023.2.2). Regions of interest included the neocortex (at the level of pallidum), midbrain (level of superior colliculi), and brainstem (mesencephalic trigeminal nucleus). Digital quantification was performed by adjusting the threshold feature in ImageJ to select positively stained pixels, and the percentage area above the threshold was quantified for analysis, blinded for the treatment group assignments. In addition, three different individuals scored the slides blindly in a semi-quantitative manner. Their ratings were normalized (Z-scored) and averaged across the three raters.

### Statistics

The data obtained in **Figure 3-5, 7-9 and Supplemental Figure 2-3, 8** were subjected to statistical analysis using a one-way analysis of variance (ANOVA) test to evaluate the differences between groups. Following the ANOVA analysis, post hoc tests were conducted using Tukey’s Honestly Significant Difference (HSD) test to determine specific group differences. For the semi-quantitative IHC data (**Figure 9C**), the non-parametric Kruskal-Wallis test was used to determine any significant group differences, with Dunn’s multiple comparisons test for the post hoc analyses to determine which groups were significantly different. The significance level was set at p < 0.05 to determine the statistical significance of the observed group differences. All statistical analyses were performed using GraphPad Prism 9.0.

**Figure 8:**
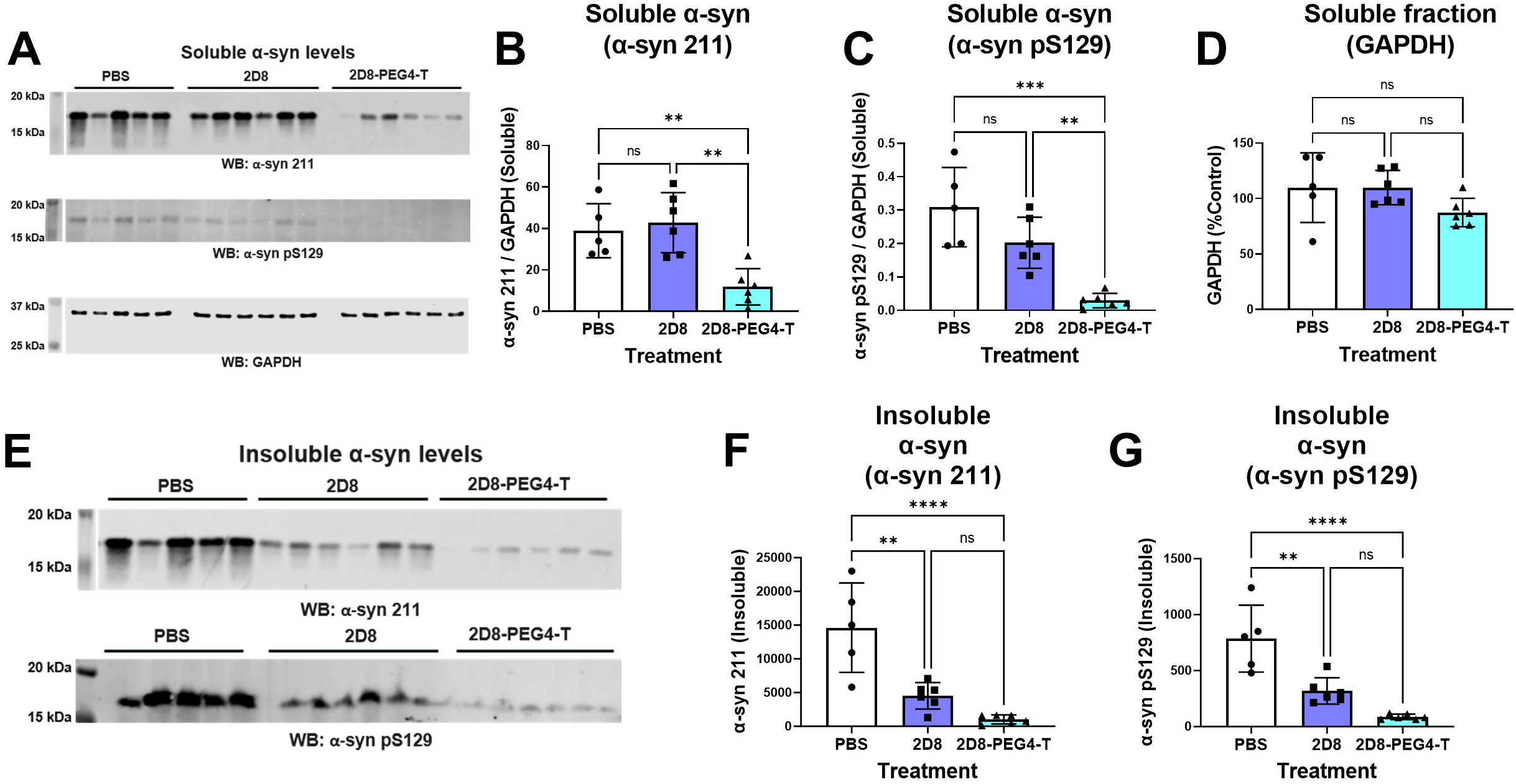
Western blot analysis of soluble and insoluble *α*-syn from the in vivo study. (A) Western blot analysis of the soluble fraction probed with total α-syn antibody 211, pS129 α-syn antibody, and loading control GAPDH antibody. The complete western blots and bands analyzed are shown in **Supplemental Figure 10**. (B) One-way ANOVA revealed group differences in soluble α-syn levels as detected by total α-syn antibody 211 (p=0.0013). sdAb 2D8-PEG4-T decreased total α-syn levels by 70% (p=0.0072), compared to the PBS treatment (p=0.0072), and by 72% compared to sdAb 2D8 (p=0.0018). (C) One-way ANOVA revealed group differences in soluble α-syn levels as detected by pS129 α-syn antibody (p=0.0002). sdAb 2D8-PEG4-T decreased pS129 α-syn levels by 90% compared to the PBS treatment (p=0.0001), and by 85% compared to sdAb 2D8 (p=0.0055). (D) One-way ANOVA did not reveal significant group differences in GAPDH levels (p=0.1369), indicating lack of toxicity of the sdAb treatments. (E) Western blot analysis showing the sarkosyl pellet insoluble fraction probed with total α-syn antibody 211 and phospho-serine129 (pS129) α-syn antibody. The complete western blots and bands analyzed are shown in **Supplemental Figure 10.** (F-G) One-way ANOVA revealed significant group differences in insoluble α-syn levels as detected by (F) total α-syn antibody 211 (p=0.0001) and (G) pS129 α-syn antibody (p<0.0001), compared to the PBS treatment. sdAb 2D8 decreased total α-syn levels by 69% (p=0.0015, F) and pS129 α-syn levels by 59% (p=0.0016, G). sdAb 2D8-PEG4-T decreased total α-syn levels by 93% (p<0.0001, F) and pS129 α-syn levels by 89% (p<0.0001, G), compared to the PBS treatment. **, ***, ****: p<0.01, 0.001, 0.0001 (Tukey’s post-hoc test).

**Figure 9:**
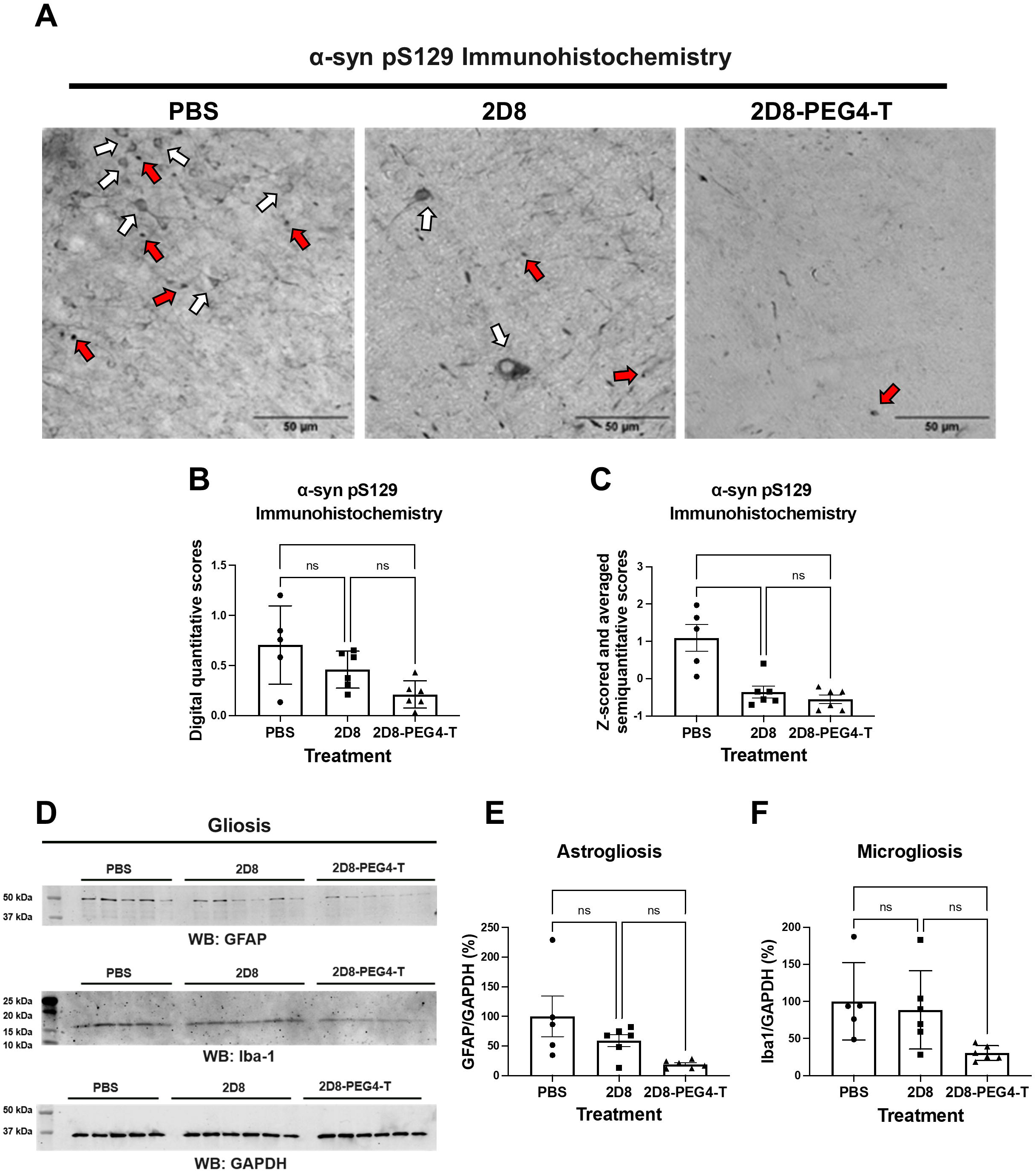
Analysis of phospho-Ser129 (pS129) immunoreactivity and astro-/microgliosis following sdAb administration in M83 mice. (A) Representative images of coronal brain sections through the midbrain region from mice treated with PBS, 2D8, and 2D8-PEG4-T, showing pS129-labeled α-syn aggregates. Distinct neuronal perikaryal inclusions (white arrows) and Lewy body (LB)-like inclusions (red arrows) were evident in the PBS treatment group. In comparison, the 2D8 treatment group exhibited reduced perikaryal inclusions (white arrows) and fewer LB-like inclusions (red arrows). Some vascular (lower half of image) and/or spheroid-like pS129 αsyn immunoreactivity (upper right of image) was also seen in this group. The 2D8-PEG4-T treatment group displayed only a few LB-like inclusions (red arrows), and some vascular immunoreactivity or perfusion artifact. Analogous α-syn pathologies were noted in the original report of this model (*51*). Scale bar: 50 μm. (B) Digital quantification of brain α-syn pathology. One-way ANOVA revealed group differences in pS129 α-syn staining (p=0.0188). 2D8-PEG4-T group had a significantly reduced α-syn pathology compared to the PBS group (70% reduction, p=0.0146). There was no significant difference between 2D8 and 2D8-PEG4-T groups. (C) Semi-quantitative analysis of brain α-syn pathology. One-way ANOVA revealed group differences in pS129 α-syn staining (p=0.0020). Both 2D8 (p=0.0487) and 2D8-PEG4-T (p=0.0079) groups had significantly reduced α-syn pathology compared to the PBS group (49% vs 62% reduction, respectively). There was no significant difference between 2D8 and 2D8-PEG4-T. (D) Western blots of the soluble brain fraction probed with GFAP antibody, Iba-1 antibody, and loading control GAPDH antibody. (E) One-way ANOVA revealed group differences in normalized GFAP levels. (p=0.028). sdAb 2D8-PEG4-T decreased GFAP levels by 81% compared to the PBS treatment (p=0.022). (F) One-way ANOVA revealed group differences in normalized Iba-1 levels. (p=0.0328). sdAb 2D8-PEG4-T decreased Iba-1 levels by 70% compared to the PBS treatment (p=0.0429). **, ***, ****: p<0.01, 0.001, 0.0001 (Tukey’s post-hoc test). The complete western blots and bands analyzed are in **Supplemental Figure 10.**

## Results

### Design and in vitro testing of targeted *α*-syn degraders

We have recently developed sdAb-based in vivo imaging probes (2D10 and 2D8) that allow for specific and non-invasive imaging of α-syn pathology in mice (*37*). The brain signal strongly correlates with lesion burden, indicating their potential as diagnostic and therapeutic candidates. These sdAbs bind well to pathological α-syn derived from human and mouse brain samples. To enhance their therapeutic potential, we conjugated the sdAbs with thalidomide, an E3 ligase ligand, to merge sdAb-based immunotherapy with PROTACs technology. Our phage display library generated from B-cells of a llama immunized with full-length rec α-syn protein yielded 58 unique anti-α-syn sdAb clones. Through iterative screening of their binding to various α-syn preparations and subsequent testing of lead sdAbs in a synucleinopathies primary culture model, we selected anti-α-syn sdAb 2D8 to determine if its PROTAC derivative would have enhanced efficacy over its unmodified version.

Specifically, we coupled sdAb 2D8 with thalidomide through three different lengths of a polyethylene glycol linker [–(PEG)n–, n = 2, 4, 6] via lysine residues of the sdAb, to optimize the orientation of the molecule to both α-syn and E3 ligase for maximum degradation efficacy (**Figure 1 and Supplemental Figure 1**). The resulting derivatives were denoted as 2D8-PEGn-Thalidomide (2D8-PEGn-T), where n represents the length of the linker (n = 2, 4, 6). The complementarity-determining regions (CDRs) of sdAbs play a crucial role in their antigen-binding specificity and affinity. As lysine residues are absent in the CDR regions of 2D8 (*58*), we assumed and verified by binding experiments that such conjugations would not affect its recognition of the α-syn protein. Specifically, we used a biolayer interferometry (BLI) assay to measure the in vitro affinity of the degrader molecule against two α-syn preparations: rec α-syn and the soluble fraction (S1) from a LBD brain, which is its most toxic fraction (**Supplemental Figure 2A-C**). Unmodified sdAb 2D8 had binding affinities of 41.2 nM and 84.1 nM against rec α-syn and the soluble S1 fraction of the human LBD brain, respectively. All the modified sdAb conjugates bound to these two α-syn preparations within the same order of magnitude as the unmodified sdAb 2D8, ranging from 41.8 to 66.7 nM against rec α-syn, and 56.2 to 107.1 nM against the soluble S1 brain fraction (**Figure 2**). Additional information on association (k_a_) and dissociation (k_d_) values can be found in **Supplemental Table 1**.

The LBD brain soluble fraction includes proteins beside α-syn. Therefore, to demonstrate the binding of unmodified and modified sdAbs to α-syn in the complex LBD brain soluble fraction, we conducted immunoprecipitation (IP) to deplete α-syn and performed dot-blot assays (see **Supplemental Figure 3A**). Initially, we conducted an IP experiment using Protein A/G beads and the α-syn 211 antibody to remove α-syn from the soluble fraction, resulting in S1 (IP α-syn flow through). We further eluted the antigen from the beads to obtain S1 (IP α-syn). Subsequent dot-blot analysis revealed significant differences in normalized dot-blot signals among the groups (p < 0.0001 for 2D8, 2D8-PEG2-T, 2D8-PEG4-T, and 2D8-PEG6-T; one-way ANOVA, see **Supplemental Figure 3B**). The results showed that both 2D8 and modified 2D8 exhibited reduced binding after α-syn depletion with binding decreasing by 90% (p < 0.0001, 2D8), 92% (p < 0.0001, 2D8-PEG2-T), 90% (p < 0.0001, 2D8-PEG4-T), and 91% (p < 0.0001, 2D8-PEG6-T) compared to S1, respectively. Notably, stronger binding of S1 (IP α-syn) was observed, with binding increasing by 86% (p < 0.0001, 2D8), 88% (p < 0.0001, 2D8-PEG2-T), 86% (p < 0.0001, 2D8-PEG4-T), and 88% (p < 0.0001, 2D8-PEG6-T) compared to S1 (IP α-syn flow through), respectively. Consequently, both 2D8 and modified 2D8 displayed binding to α-syn in the LBD S1 solution, indicating their potential as therapeutic candidates targeting α-syn in this soluble fraction. And these results confirm that conjugating thalidomide via linkers of three different lengths does not affect α-syn binding of the sdAb 2D8.

### Degradation efficacy of sdAbs targeting *α*-syn in a primary neuronal culture model of synucleinopathy

As previously discussed, we proposed that antibody therapy can target proteinopathies both intra- and extracellularly, contingent on the antibodies’ ability to enter neurons (*27, 38*). We have modeled these two mechanisms in primary cultures using human derived PHF from tauopathy patients for anti-tau antibody immunotherapy studies (*39, 40, 59*). A similar approach is applicable to synucleinopathies. Various in vitro and animal models have established that pathological α-syn seeds generated from diseased brains can accelerate the development of synucleinopathy (*60–63*). To establish such culture model, various α-syn fractions derived from human LBD brains were prepared (**Supplemental Figure 2A**). Subsequently, their toxicity was evaluated using a primary mouse neuronal culture model. The findings revealed that the soluble S1 fraction was most toxic and significantly seeded α-syn in the neuronal culture model. Soluble S1 fraction induced significant toxicity, with reduced GAPDH levels by 67% (p = 0.0005) and 65% (p = 0.0001) on day 5 and day 7, compared to the control group, respectively. (**Supplemental Figure 2B-C**). We further accessed the seeding capacity of the soluble S1 fraction. To achieve that, primary neuronal cultures were established from M83 mouse pups. These cultures were exposed to 10 μg/ml of the soluble S1 fraction for varying durations (1, 2, 3, 5 and 7 days). Subsequently, to assess the seeding capacity of the soluble S1 fraction, we conducted dot-blot assays employing conformational antibodies A11 and OC, which were demonstrated to detect oligomeric amorphous and fibrillary aggregates, respectively (*52, 53*). Our findings, detailed in **Supplemental Figure 4**, revealed the clear formation of α-syn aggregates on days 3, 5, and 7 in the presence of the soluble S1 fraction, confirming the seeding capacity of the LBD brain soluble S1 fraction. Quantitative analysis of the dot blot signals demonstrated significantly elevated levels of α-syn aggregates in the presence of the LBD brain soluble S1 fraction (A11 antibody: p = 0.0241 (day 3), p = 0.0048 (day 5), p = 0.0151 (day 7); OC antibody: p = 0.0007 (day 3), p = 0.0286 (day 5); unpaired t test). These results provide evidence supporting the toxicity and seeding capacity of the S1 LBD brain fraction evaluated in our study. Consequently, we incubated primary transgenic M83 A53T α-syn neurons with LBD brain soluble S1 α-syn fraction and treated them with sdAbs using the (S1 + sdAb) or (S1 → sdAb) dosing paradigms, which reflect extracellular vs intracellular interaction of the sdAb with the α-syn target, respectively (**Figures 3A and 4A**).

To determine extracellular clearance, primary neurons were incubated with 10 µg/ml of LBD brain-derived soluble S1 α-syn fraction and treated with 5 μg/ml of 2D8, or its molar equivalent of modified 2D8, simultaneously for 24 h. Following this, the media was replaced, the cells were maintained in culture for an additional 96 h, after which samples were collected for analyses (**Figure 3A**). The results indicated that groups differed significantly in their GAPDH levels (p < 0.0001, one-way ANOVA, **Figure 3B and D**). S1 alone induced significant toxicity (57% of controls, p = 0.0003), whereas both unmodified sdAb 2D8 and the three modified sdAb 2D8-PEGn-T (n = 2, 4, 6) prevented S1-induced toxicity, with increased GAPDH levels by 42% (2D8, p = 0.0005), 43% (2D8-PEG2-T, p = 0.0003), 52% (2D8-PEG4-T, p < 0.0001) and 39% (2D8-PEG6-T, p = 0.0018), compared to that of S1 alone, respectively. Apart from the GAPDH marker, we also utilized NeuN marker to further evaluate the toxicity of pathological α-syn in neuronal cultures (**Figure 3C**). The results align closely with the findings from GAPDH levels presented in **Figure 3B**, demonstrating significant differences in NeuN levels among the groups (p < 0.0001, one-way ANOVA, **Figure 3C and E**). Notably, S1 alone induced substantial toxicity (46% of controls, p < 0.0001), while both unmodified sdAb 2D8 and the three modified sdAbs 2D8-PEGn-T (n = 2, 4, 6) mitigated S1-induced toxicity, leading to increased NeuN levels by 50% (2D8, p < 0.0001), 45% (2D8-PEG2-T, p = 0.0017), 52% (2D8-PEG4-T, p < 0.0001), and 52% (2D8-PEG6-T, p < 0.0001) compared to S1 alone, respectively. Our findings are consistent with our previous study where both NeuN and GAPDH yielded similar toxicity results when using anti-tau sdAb to block tau toxicity and enhance its clearance (*39*).

In terms of the α-syn levels, one-way ANOVA revealed a significant group difference in normalized α-syn levels (α-syn/GAPDH, p < 0.0001, **Figure 3D-E**). S1 alone increased intracellular α-syn levels by 39% compared to control values at 96 h (p = 0.0002), while both unmodified sdAb 2D8 and the three modified sdAb 2D8-PEGn-T (n = 2, 4, 6) prevented α-syn seeding This resulted in lower intracellular α-syn levels relative to S1 alone (decreased total α-syn/GAPDH levels by 35% (2D8, p = 0.0008), 45% (2D8-PEG2-T, p < 0.0001), 48% (2D8-PEG4-T, p<0.0001) and 47% (2D8-PEG6-T, p < 0.0001), compared to S1 alone values, respectively.

Notably, in this extracellular blockage dosing paradigm, there was no significant difference between the unmodified sdAb 2D8 and the three modified sdAb 2D8-PEGn-T (n=2, 4, 6) in terms of GAPDH and normalized α-syn levels. Outside of the cell, sdAbs presumably sequester α-syn aggregates and interfere with their assembly and/or neuronal uptake, collectively preventing the spread and seeding of α-syn pathology (*27*). These results were in line with the binding experiment (**Figure 2**), which indicated that the chemical modification did not affect the affinity of the sdAb for α-syn.

Concurrently, intracellular clearance was assessed by incubating primary M83 transgenic neurons with 10 µg/ml LBD brain-derived soluble fraction α-syn (S1) for 24 h, which allowed the cells to take up the pathological α-syn. The media was then exchanged, and various concentrations of sdAbs were added and incubated for an additional 4 days (S1 → sdAb). Because the brain S1 α-syn fraction is already internalized in this paradigm, the sdAbs must also enter the cells to be effective. PROTACs in general can show a Hook Effect phenomenon at high concentration (*64*), which may be due to the formation of unproductive PROTAC dimers, rather than the productive trimeric complex required for degradation (**Figure 4B**). Therefore, five different concentrations were tested of 2D8, ranging from 0.625-10 µg/ml, or its molar equivalent of modified 2D8.

One-way ANOVA revealed significant differences in normalized α-syn levels when the concentration of sdAbs was higher than 0.625 µg/ml (p = 0.02 for 1.25 µg/ml, p = 0.0004 for 2.5 µg/ml, p = 0.0001 for 5.0 µg/ml, p = 0.0136 for 10.0 µg/ml, **Figure 4C-D**). The S1 α-syn fraction alone led to a 46% increase in intracellular α-syn levels, compared to control values at 96 h (1.25 µg/ml graph, p = 0.0154; 2.5 µg/ml graph, p = 0.0049; 5.0 µg/ml graph, p = 0.0043). At 2.5-5 µg/ml, 2D8-PEG4-T significantly prevented the S1-induced increase in α-syn levels (reduced by 61% at 2.5 µg/ml (p = 0.0001) and 67% at 5.0 µg/ml (p < 0.0001), relative to S1 alone). Compared to the unmodified sdAb 2D8, sdAb 2D8-PEG4-T significantly reduced α-syn levels at 2.5 and 5 µg/ml (38% reduction at 2.5 µg/ml (p = 0.0298) and 41% reduction at 5.0 µg/ml (p = 0.0126), relative to unmodified sdAb 2D8). Therefore, we concluded that modified sdAb 2D8 with four PEG linkers exhibited a stronger degradation capacity than unmodified 2D8 when the concentration ranged from 2.5 to 5 µg/ml. However, this effect may saturate at higher concentration like 10 µg/ml, at which dose group differences were not as pronounced. Based on these results, we selected modified 2D8 with four PEG linkers as a candidate for subsequent studies.

As previously described, we conducted BLI binding experiments to confirm that the modified sdAb 2D8 retained the affinity of its unmodified version for α-syn preparations (**Figure 2**). Our previous work showed that unmodified sdAb 2D8 readily entered the brain of M83 synucleinopathy mice and bound to pathological α-syn protein intracellularly after a single intravenous (i.v.) injection (*37*). To determine whether 2D8-PEG4-T could bind to α-syn protein intracellularly in a similar manner as its unmodified version, primary M83 transgenic neurons were incubated with 10 µg/ml of LBD brain-derived soluble S1 α-syn fraction for 24 h. The culture was then washed and treated with 5 µg/ml of either unmodified sdAb 2D8 or modified sdAb 2D8-PEG4-T for 2 h, followed by immunohistochemistry and analysis by confocal microscopy (**Figure 4E**). The results showed that both unmodified sdAb 2D8 and modified sdAb 2D8-PEG4-T were taken up by the neurons and co-localized with α-syn protein (Mander’s coefficients: M2: 0.481 (ratio of α-syn colocalized with 2D8); M2: 0.445 (ratio of α-syn colocalized with 2D8-PEG4-T)). This indicates that both versions of the 2D8 sdAb bind to intracellular α-syn to a similar degree.

### Degradation pathways of sdAbs targeting *α*-syn in a primary neuronal culture model of synucleinopathy

The degradation mechanism of 2D8 and 2D8-PEG4-T was investigated by identifying which of the two major degradation pathways was likely to be involved (*27*), namely the lysosomal and proteasomal pathways (**Figure 5A**). To achieve this, lysosome inhibitor bafilomycin A1 (Baf.A1) or proteasome inhibitor MG132 were used to block these two pathways. Specifically, M83 primary neurons were incubated with 10 µg/ml of LBD brain-derived S1 soluble α-syn fraction for 24 h to allow the pathological α-syn to be taken up by the cells. Then, the media was exchanged, and the neurons were pre-treated for 6 h with either lysosome inhibitor Baf.A1 or proteasome inhibitor MG132 followed by 2 days treatment with 5 µg/ml sdAbs (**Figure 5B**). Similar to our prior results (see **Figure 4C-D**), S1 alone increased normalized intracellular α-syn levels by 59% in the Baf.A1 study (**Figure 5C-D**), compared to control values (p = 0.0013 for **Figure 5C**; p < 0.0001 for **Figure 5D**, relative to the control group). Likewise, S1 alone increased intracellular α-syn levels by 53% in the MG132 study (**Figure 5E-F**), compared to control values (p = 0.0077 for **Figure 5E**; p = 0.0018 for **Figure 5F**, relative to the control group). Both unmodified sdAb 2D8 and modified sdAb 2D8-PEG4-T significantly prevented the S1-induced increase in α-syn levels in the Baf.A1 study (40% reduction, p = 0.0194 for 2D8, **Figure 5C**; and 83% reduction, p < 0.0001 for 2D8-PEG4-T, **Figure 5D**, relative to S1 alone). However, this α-syn clearance was fully blocked by the Baf.A1 inhibitor for 2D8 but only partially blocked for 2D8-PEG4-T. Specifically, α-syn levels were increased by 34% for 2D8 with 0.5 µM Baf.A1 inhibitor, relative to 2D8 alone (p = 0.0494, **Figure 5C**), and increased by 22% for 2D8-PEG4-T with 1 µM Baf.A1 inhibitor, relative to 2D8-PEG4-T alone (p = 0.0054, **Figure 5D**). Additionally, both unmodified sdAb 2D8 and modified sdAb 2D8-PEG4-T significantly prevented the S1-induced increase in α-syn levels in the MG132 study (40% reduction, p = 0.0434 for 2D8, **Figure 5E**; and 76% reduction, p < 0.0001 for 2D8-PEG4-T, **Figure 5F**; relative to S1 alone). The MG132 proteasomal inhibitor did not affect the efficacy of 2D8 (**Figure 5E**), but it decreased the efficacy of 2D8-PEG4-T in a concentration-dependent manner (**Figure 5F**, α-syn levels were increased by 32% with 5 µM MG132 (p = 0.0437) and increased by 48% with 10 µM MG132 (p = 0.004), relative to 2D8-PEG4-T).

In summary, the results indicate that unmodified sdAb 2D8 predominantly induces α-syn degradation through lysosomal pathways, whereas the modified sdAb 2D8-PEG4-T is capable of both utilizing this pathway and activating the previously untapped proteasomal degradation pathway, leading to superior degradation efficacy compared to sdAb 2D8.

Next, we conducted co-immunoprecipitation (co-IP) assays in M83 primary neurons treated with sdAb to investigate the ternary complex interactions predicted for α-syn (**Figure 6A**). M83 primary neurons were exposed to 10 µg/ml of LBD brain-derived soluble S1 α-syn fraction for 24 h to allow it to be sufficiently taken up by the cells. Next, the media was changed, and the neurons were pre-treated with the proteasome inhibitor MG132 (10 µM) for 6 h to promote ternary complex accumulation before α-syn clearance. Subsequently, 5 µg/ml of unmodified sdAb 2D8 or modified sdAb 2D8-PEG4-T were directly added to the media without exchanging it. Samples were collected 2 days after antibody application **(Figure 6B).** Following the co-IP experiment, we confirmed the successful capture of the α-syn protein by incubating with α-syn antibody (PA5-13397, **Figure 6C**). After conducting the analysis, we found significant differences between groups in long conjugations to α-syn (**Figure 6C (i), Supplemental Figure 8A,** one-way ANOVA, p = 0.0017). Notably, treating modified sdAb 2D8-PEG4-T with the proteasome inhibitor MG132 resulted in a 56% increase in long conjugations to α-syn compared to untreated sdAb 2D8 with MG132 (p = 0.0245). These elongated conjugations were identified as ubiquitin chains via detection with a ubiquitin antibody (P4D1, **Figure 6D (i)**). Additionally, we observed significant group differences in ubiquitin levels (**Figure 6D (i), Supplemental Figure 8B**, one-way ANOVA, p < 0.0001). Remarkably, modified sdAb 2D8-PEG4-T significantly elevated ubiquitin levels by 53% (p = 0.0064) and 43% (p = 0.0208) compared to S1 alone and sdAb 2D8, respectively. Treatment with the proteasome inhibitor MG132 notably increased ubiquitin levels by 42% (p = 0.027, sdAb 2D8 vs sdAb 2D8 + MG132) and 49% (p = 0.0002, sdAb 2D8-PEG4-T vs sdAb 2D8-PEG4-T + MG132) for both unmodified and modified sdAb 2D8 groups. Furthermore, modified sdAb 2D8-PEG4-T combined with MG132 led to a 50% increase in ubiquitin levels compared to untreated sdAb 2D8 with MG132 (p = 0.0001).

Ubiquitin levels were low in control and S1-alone treatment groups, suggesting basal interaction between α-syn and ubiquitin in primary neurons, consistent with findings in mouse brain tissue (*65*). Unmodified sdAb 2D8 treatment did not increase ubiquitin levels, although MG132 slightly increased it. In contrast, modified sdAb 2D8-PEG4-T treatment increased long ubiquitin chain conjugation on α-syn, and MG132 further increased ubiquitin levels (**Figure 6D** (i)). The degradation effect of modified 2D8-PEG4-T was confirmed by IP input-only western blotting analysis of total protein lysate without subjecting it to immunoprecipitation. Additionally, treatment with the proteasome inhibitor MG132 prevented the observed degradation effect (**Figure 6E**). Collectively, these results support ternary complex formation between α-syn, 2D8-PEG4-T, and E3 ligase, followed by increased α-syn ubiquitination and proteasomal degradation.

### Degradation efficacy of sdAbs targeting *α*-syn in an M83 mouse model of synucleinopathy

In the next phase of our study, we assessed the in vivo effectiveness of the unmodified and modified sdAb in an M83 mouse model of synucleinopathy. Previous findings from our diagnostic imaging study indicated that the anti-α-syn sdAbs 2D8 and 2D10 traversed the blood-brain barrier and colocalized with intraneuronal α-syn aggregates after an intravenous injection in this mouse model (*37*). The intensity of the sdAb brain signal correlated strongly with the levels of pathological α-syn in the brain, with 2D10 showing a clear superiority over 2D8 in signal intensity, and a slight advantage in degree of correlation and specificity for α-syn lesions. However, in our pilot culture studies, PROTAC-modified 2D10 was less efficacious than PROTAC-modified 2D8. This discrepancy may be attributed to potential instability of the overall conformation of α-syn, PROTAC-modified 2D10, and E3 ligase, which could then prevent generation of the ternary complex (**Figure 4B**) (*64*). Its formation is crucial for the efficacy of ubiquitination, and thereby proteasomal degradation. These prior findings support 2D10’s potential as a diagnostic imaging probe, and modified 2D8’s potential as a therapy.

Taking advantage of this prior work, we initially evaluated on Day 1 α-syn aggregate brain burden in 7-8-month-old homozygous M83 mice (n = 17) by IVIS imaging following a single intravenous injection of near-infrared labeled 2D10 sdAb (**Figure 7A-C**). The mice should have moderate to severe pathological α-syn burden at that age. Their individual brain signal allowed us to split them into three different treatment groups with a comparable average brain signal and thereby presumably similar α-syn brain burden (n = 5-6 per group, **Figure 7B-C**). The mice then received i.v. injections on Day 4, 7 and 10 of PBS (controls), unmodified sdAb 2D8, or modified sdAb 2D8-PEG4-T in PBS, respectively. Subsequently, all mice were injected with the near-infrared labeled 2D10 to re-evaluate α-syn pathology. Afterwards, the mice were perfused, and their brains extracted for tissue analyses. Significant group differences were detected in cumulative α-syn brain signal after the acute treatment period (**Figure 7B and D**, one-way ANOVA, p = 0.0059). Remarkably, the administration of modified sdAb 2D8-PEG4-T led to an 81% reduction in α-syn brain signal compared to the PBS control group (p = 0.0049) (**Figure 7 B and D**). There was a trend for decreased α-syn brain signal in the 2D8 group, but it did not differ significantly from the control or the 2D8-PEG4-T group.

Following the imaging studies, we homogenized the left-brain hemisphere of the mice and prepared both soluble and insoluble fractions of α-syn protein. GAPDH levels did not differ between the groups (**Figure 8A and D**, one-way ANOVA, p = 0.1369), indicating absence of treatment toxicity. For soluble α-syn, only modified sdAb 2D8 significantly reduced soluble total (70%, p = 0.0072) and phospho-Ser129 α-syn (90%, p = 0.0001), compared to controls. Moreover, it exhibited significant clearance of both soluble total and phospho-Ser129 α-syn, compared to the unmodified sdAb 2D8 treatment (p < 0.01). (**Figure 8A-D)** Total and phospho-Ser129 insoluble α-syn differed significantly between the treatment groups (**Figure 8E-G**, one-way ANOVA, p = 0.0001 (total insoluble α-syn), p < 0.0001 (phospho-Ser129 insoluble α-syn)). Unmodified sdAb 2D8 reduced total (69%, p = 0.0015) and phospho-Ser129 (59%, p = 0.0016) insoluble α-syn, compared to the PBS control group. Likewise, modified sdAb 2D8 reduced total (93%, p < 0.0001) and phospho-Ser129 (89%, p < 0.0001) insoluble α-syn, compared to the PBS control group.

Subsequently, we conducted immunohistochemistry analysis on the right-brain hemisphere of mice using the phospho-Ser129 antibody (*66–68*), as detailed in the Methods section. Within the group of mice treated with PBS, we observed phosphorylated α-syn aggregates in various brain regions, including the neocortex, midbrain, and brainstem (**Figure 9A**). Notably, acute treatment with both 2D8 and 2D8-PEG4-T reduced these aggregates when compared to the control group (**Figure 9A**) as determined by quantitative and semi-quantitative analysis (**Figure 9B-C**). The quantitative analysis revealed significant group differences (one-way ANOVA, p = 0.0188), with 2D8-PEG4-T decreasing phospho-Ser129 immunoreactivity by 70% (p = 0.0146). Likewise, the semi-quantitative analysis revealed significant group differences (one-way ANOVA, p = 0.0020), with 2D8 and 2D8-PEG4-T decreasing phospho-Ser129 staining by 49% (p=0.0487) and 62% (p = 0.0079), respectively, compared to control group. The two treatment groups did not differ significantly in either analysis.

It is worth noting that the majority of phospho-Ser129 α-syn species are typically found in the insoluble brain fractions (*69*), and insoluble α-syn levels did not differ significantly between the 2D8 and 2D8-PEG4-T treatment groups on western blots (**Figure 8E-G**), which fits the immunohistochemical analyses. Because insoluble proteins primarily undergo lysosomal degradation (*70*), these findings indicate that unmodified sdAb 2D8 primarily induces α-syn degradation through the lysosomal pathway, while the modified sdAb 2D8-PEG4-T works both via this pathway in addition to enhancing proteasomal clearance of soluble α-syn, resulting in superior degradation efficacy compared to 2D8. This is supported both by the culture (**Figures 4C-D, 5C-H**) and in vivo studies (**Figure 8A-G**).

The pathogenesis of synucleinopathy is closely linked to an intense glial response (*71–73*). To assess whether 2D8 and 2D8-PEG4-T can mitigate these immune reactions within glial cells, we examined the levels of GFAP and Iba1 levels of the mice brain after the treatment, as markers for astrocytes and microglia. To this end, we performed western assays using the low-speed supernatant (LSS) of the mouse brain (see method section). Normalized GFAP (**Figure 9D-E)** and Iba-1 (**Figure 9D, and F)** levels differed significantly between the treatment groups (one-way ANOVA, p = 0.028 (GFAP), p = 0.033 (Iba-1)). Specifically, 2D8-PEG4-T reduced GFAP by 81% (p = 0.022) and Iba-1 by 70% (p = 0.043), compared to the PBS control group. As for insoluble α-syn, 2D8 and 2D8-PEG4-T groups did not differ significantly in their effects on GFAP and Iba-1 levels. Overall, these findings suggest that the 2D8-PEG4-T mediated clearance of α-syn alleviates gliosis within the brain.

Overall, these results provide insight into pathways involved in sdAb-mediated clearance of pathological α-syn, support therapeutic candidate development of the 2D8 sdAb and show the utility of in vivo sdAb-mediated imaging of α-syn for treatment group assignments and to monitor treatment efficacy.

## Discussion

We have successfully improved the efficacy of an sdAb targeting α-syn for clearance by enhancing its proteasomal degradation capacity while maintaining its lysosomal clearance ability and affinity for recombinant and human LBD brain derived α-syn. This dual-targeting approach significantly enhanced the clearance of α-syn in primary culture and mouse models of synucleinopathy and was associated with reduced astro- and microgliosis in the mice. Remarkably, only three i.v. injections within a week reduced α-syn levels in the brain by 70-93%. The in vivo study was facilitated by our ability to assess by imaging in intact animals their α-syn brain burden prior to and post-treatment, with the latter imaging data corresponding well with biochemical and histological analyses of their α-syn brain pathology. These findings highlight the potential of our sdAb-based protein degrader as a promising therapy to clear pathological α-syn, and of the related in vivo α-syn imaging approach to assign animals into treatment groups with similar α-syn burden as well as to monitor the efficacy of therapies.

To evaluate the degradation efficacy of the sdAbs, we employed a culture model of synucleinopathy using transgenic M83 primary neurons with A53T α-syn mutation combined with LBD brain soluble S1 α-syn fraction. While the majority of LBD patients do not possess the A53T mutation, this mutation has been found to increase the self-aggregation rate of the protein, leading to accelerated synucleinopathy progression in relevant mouse models (*51, 74*). Consequently, the rationale behind utilizing primary neuronal cultures derived from M83 mice carrying the A53T α-syn mutation, along with the addition of LBD brain soluble fraction, was based on the aim to expedite disease propagation through a prion-like mechanism (*75–77*). It is worth noting that the binding epitope of 2D8, amino acids 71-94, is well outside the A53T mutation site (*78*).

We developed the sdAb-based protein degrader by coupling sdAb 2D8 with thalidomide through PEG linker of different lengths via lysine residues of the sdAb. The optimal design of the PEG linker length is paramount for modulating the physicochemical properties and spatial orientation of the ternary complex (sdAb-α-syn-E3 ubiquitin ligase), ultimately influencing the efficiency of target protein degradation. We employed three different conformationally flexible PEG linkers to strategically position the sdAb for simultaneous binding to both the E3 ligase and the α-syn protein. This strategic design aimed to promote the formation of a stable trimeric complex (**Figure 3-4**), facilitating subsequent ubiquitination of α-syn and its proteasomal degradation.

Furthermore, the selection of thalidomide as an E3 ligase recruiter, specifically targeting CRBN, stems from its ability to effectively engage the E3-ubiquitin ligase CRL4^CRBN^ (*79*). Thalidomide and its derivatives have shown promising results in inducing the degradation of various target proteins, highlighting their potential as a versatile moiety for PROTAC-mediated protein degradation strategies (*79–84*). Notably, CRBN has been successfully recruited in a different PROTACs strategy to specifically degrade the α-syn protein (*85*). Interestingly, compared to another E3 ligase Von Hippel-Lindau (VHL) (*86–88*), CRBN may have a broader acceptance of neosubstrate degradation making it a more favorable candidate for α-syn protein degradation in our study (*89*).

Regarding the precise attachment point within the sdAb, we opted to attach thalidomide to the lysine residues of the sdAb. Given the critical role of the CDRs in sdAb antigen-binding specificity and affinity, careful consideration was given to not use sdAb with lysine residues within these pivotal regions to preserve the binding characteristics of the sdAb.

In our studies, we have shown that antibody therapy can effectively target proteinopathies both intra- and extracellularly by gaining access to neurons, for reviews see (*27, 90*). We successfully modeled these mechanisms in primary cultures by introducing pathological α-syn seeds derived from diseased brains. It is worth noting that while some internalization of sdAb and LBD S1 solution α-syn may occur before medium exchange in the extracellular mechanism, the predominant mode of interaction is likely extracellular. Similarly, in the intracellular mechanism, a minor fraction of brain S1 α-syn on the membrane may not be completely removed by washing and medium exchange, but the majority of the interaction is anticipated to be intracellular. To confirm this, we previously showed that the uptake of antibodies into neurons can be inhibited by an antibody targeting FcII/III receptors or by using dansyl cadaverine, which hinders receptor-mediated endocytosis (*91*). In co-incubation scenarios where the sdAb and pathological protein were introduced simultaneously, blocking antibody uptake did not impact the results. However, introducing the antibody 24 hours after the addition of pathological proteins and blocking its uptake hindered its beneficial effects. These results affirm that the antibody operates extracellularly during co-incubation but exerts intracellular effects when added 24 hours after pathological protein has been added to the culture (*40*).

To enhance α-syn degradation through the proteasome pathway, modifications have been introduced to antibody fragments by fusing highly negatively charged proteasomal retargeting sequences (PEST motifs), resulting in enhanced degradation of GFP-labeled α-syn protein when expressed together in St14A cells (*92, 93*). Moreover, this intrabody approach has been examined in vivo using gene therapy delivery of two proteasome-directed sdAbs in a synucleinopathy rat model (*94*). By performing unilateral substantia nigra injections of vectors carrying the modified sdAbs, a reduction in phospho-Ser129 α-syn labeling was observed compared to saline-treated animals. However, it remains unclear whether soluble or insoluble α-syn was reduced, and the study did not investigate the involvement of the lysosome pathway. In addition, it has not been reported if these sdAbs are efficacious when administered directly like in our study, instead of using transfection or gene therapy approaches. Direct injection has generally not been considered feasible for therapy with sdAbs because of their short half-life but our findings clearly show the therapeutic potential of their i.v. route of administration, in particular as a PROTAC derivative.

Insoluble α-syn is more likely to undergo degradation in the lysosome due to its limited entry into the proteasome compartment (*70*). As shown in our culture studies, both unmodified sdAb 2D8 and modified sdAb 2D8-PEG4-T promote the degradation of α-syn through the lysosomal pathway (**Figure 5C-D**). Furthermore, our previous findings demonstrated that following its i.v. injection, near-infrared-labeled sdAb 2D8 enters the brain and neuronal cells, where it colocalizes with pathological α-syn within the endosome-lysosome system (*37*). In addition, the in vivo studies show that acute treatment with unmodified 2D8 clears insoluble α-syn but not soluble α-syn, which supports its efficacy via lysosomal clearance (**Figure 8**). On the other hand, the modified 2D8 sdAb works via both the lysosomal pathway and the proteasomal pathway, significantly reducing both insoluble and soluble α-syn (**Figure 5D and F**, **Figure 6, and Figure 8**). Its ability to enhance proteasomal degradation of α-syn, in addition to sustaining its lysosomal clearance, likely explains its superior efficacy over the unmodified 2D8. Notably, group differences in the post-treatment brain α-syn imaging signal in intact animals (**Figure 7D**) corresponded to biochemical and histological analysis of α-syn, in particular to soluble phospho-Ser129 α-syn, insoluble total and phospho-Ser129 α-syn on western blots of brain homogenate fractions, and phospho-Ser129 α-syn immunohistochemistry of brain sections (**Figure 8A-G**, **Figure 9A-C**).

It is worth noting that current therapies targeting α-syn primarily rely on whole IgG antibodies, which have limited brain penetrance that may diminish their effectiveness. In contrast, sdAbs because of their smaller size (15 kDa vs 150 kDa), enter the brain more readily, diffuse better into tissue, and can bind to cryptic epitopes not previously recognized by whole antibodies. Similar strategy may apply to immunotherapies targeting other neurodegenerative disease. For example, lessons from anti-amyloid-β (Aβ) antibody therapies like lecanemab (*95*) and donanemab (*96*) can be instructive. These antibodies have demonstrated a modest yet significant impact on slowing cognitive decline in phase III trials. However, it is important to note that some adverse events have been observed, such as extensive vascular inflammation and ruptured blood vessels leading to brain bleeding (*97*). One potential explanation for these events could be that anti-Aβ antibodies bind to vascular Aβ deposits, triggering the complement cascade and leading to the formation of the membrane attack complex, which in turn damages blood vessels (*98*). Additionally, complement proteins C3a and C5a may activate microglia in the vicinity, causing vascular inflammation (*99, 100*). Both lecanemab and donanemab are humanized IgG1 antibodies (*95, 96*), known for their strong effector functions that can activate various immune responses and effector mechanisms (*101*). To address safety and efficacy concerns, sdAbs present a promising alternative due to their smaller size, which can reduce immunogenicity and enhance tissue penetration compared to traditional antibodies with Fc regions. Importantly, sdAbs lack the Fc region, thereby eliminating the potential for Fc-mediated effector functions like antibody-dependent cellular cytotoxicity (ADCC) and complement-dependent cytotoxicity (CDC), which can contribute to adverse effects (*102*).

Intriguingly, recent research has found that the engagement of the E3 ubiquitin ligase TRIM21 is crucial for enabling antibodies to clear tau seeds (*103*). That particular study showed that a tau-clearing antibody became ineffective in tauopathy mice lacking TRIM21. Cytosolic TRIM21 binds to full-size antibodies via their Fc domain, stimulating proteasomal degradation of the tau-antibody complex. That finding strengthens the notion that antibodies exert their action inside neurons to eliminate aggregated tau, as we and others have previously noted in several studies (*27, 38–45*), and a similar mechanism may apply to synucleinopathies.

Consequently, the advantages provided by the smaller size of sdAbs may be counteracted by their lack of Fc domain. In addition, their small size allows for their rapid elimination from the bloodstream, as they fall well below the renal clearance threshold of approximately 30-50 kDa. Therefore, the half-life of sdAbs in blood is relatively short, typically less than 2 h, rendering them potentially less suitable as therapeutic candidates compared to whole antibodies that usually have blood half-life of 1-3 weeks (*104*).

However, during our imaging study, we observed that at the time point of 24 and 96 hours following a single intravenous injection of near-infrared-labeled 2D8, its brain signal registered at 47% and 22% of peak signal, respectively (**Supplemental Figure 9**, (*37*)). The relatively slow brain clearance can be attributed to the binding of sdAbs to α-syn aggregates within neurons. Our binding results demonstrate that both 2D8 and modified 2D8-PEG4-T sdAbs exhibit high binding affinity for α-syn, ranging from 41.2 to 85.9 nM (**Figure 2**). High binding affinity can result in a stable complex that takes longer to dissociate and be cleared from the brain (**Supplemental Figure 9**). The binding curve (**Figure 2**) and dissociation rate (**Supplemental Table 1**) indicate that once sdAbs bind to α-syn, they require time to dissociate from the antigen.

Therefore, unmodified sdAbs may have therapeutic potential despite rapid blood clearance in light of their target engagement within the brain, as supported by our in vivo efficacy findings (**Figure 8**). However, to counteract these deficiencies, modified sdAbs such as our 2D8-PEG4-T with enhanced degradation capabilities, is likely to be a superior therapeutic approach over unmodified sdAb, unless perhaps when given as a gene therapy with continuous expression.

It is important to acknowledge the limitations of our study, which primarily assessed the therapeutic efficacy of the sdAb-based protein degrader in preclinical rodent models. Further research is required to evaluate its safety, efficacy, and pharmacokinetic profile in human subjects. Additionally, the long-term effects and potential side effects associated with promoting proteasomal degradation of α-syn should be carefully investigated, although toxicity was not observed in our primary culture or mouse models of synucleinopathy.

## Conclusion

In conclusion, our study demonstrates the potential of sdAb-based protein degraders as a promising therapeutic strategy for synucleinopathies. These innovative degraders may provide new hope for patients suffering from these diseases by addressing the underlying pathology and offering a potential disease-modifying treatment option.

## List of abbreviations

α-syn: α-synuclein
SdAb: Single-domain antibody
CRBN: Cereblon
PD: Parkinson’s disease
LBD: Lewy body dementia
MSA: Multiple System Atrophy
ScFv: Single-chain variable fragments
AAV: Adeno-associated virus
Rec: recombinant
PROTACs: PROteolysis TArgeting Chimeras
POI: Protein of interest
2D8-PEG4-T: 2D8-PEG4-Thalidomide
BLI: Bio-layer interferometry
Tris-NTA: Tris-nitrilotriacetic
TBS: Tris buffered saline
IP: Immunoprecipitation
Co-IP: Co-immunoprecipitation
CDRs: Complementarity-determining regions
ADCC: Antibody-dependent cellular cytotoxicity
CDC: Complement-dependent cytotoxicity
IVIS: In Vivo Imaging System
ROI: Region of interest
LSS: Low-Speed Supernatant
SP fraction: Sarkosyl pellet fraction

## Declarations

### Ethics approval

Any data derived from human tissue would not include any patient identifiers. This study was conducted in accordance with US ethical guidelines and was deemed exempt from ethics approval by the Institutional Review Board (IRB). All mouse experiments were performed under an institutional animal care and use committee (IACUC) approved protocol with the mice housed in Association for Assessment and Accreditation of Laboratory Animal Care (AAALAC) approved facilities with access to food and water ad libitum.

### Consent for publication

Not applicable

### Availability of data and materials

Data and materials availability: All data needed to evaluate the conclusions in the paper are present in the paper and/or the Supplemental Materials.

## Authors’ contributions

Y.J. performed most of the studies and related analyses with help from C.J. for the immunofluorescence assay, Y.L. and A. M. T. for the sectioning and immunohistochemistry staining of the mouse brains and related analyses. Y. L. maintained the animal colonies. E.E.C. generated the enriched α-syn fractions from the LBD brain and compared their initial toxicity in culture. R.P. and X.-P.K. helped with mammalian expression and purification of the 2D8 and 2D10 sdAb. Y.J. and E.M.S. designed the experiments and wrote the article. All authors had the opportunity to edit the article. E.M.S. supervised the project.

## Acknowledgements

We are grateful to the National Disease Research Interchange (NDRI, Philadelphia, PA), from which we purchased a brain with extensive cortical α-syn inclusions, which is a hallmark of LBD.

---

**Supplemental Figure 1:**
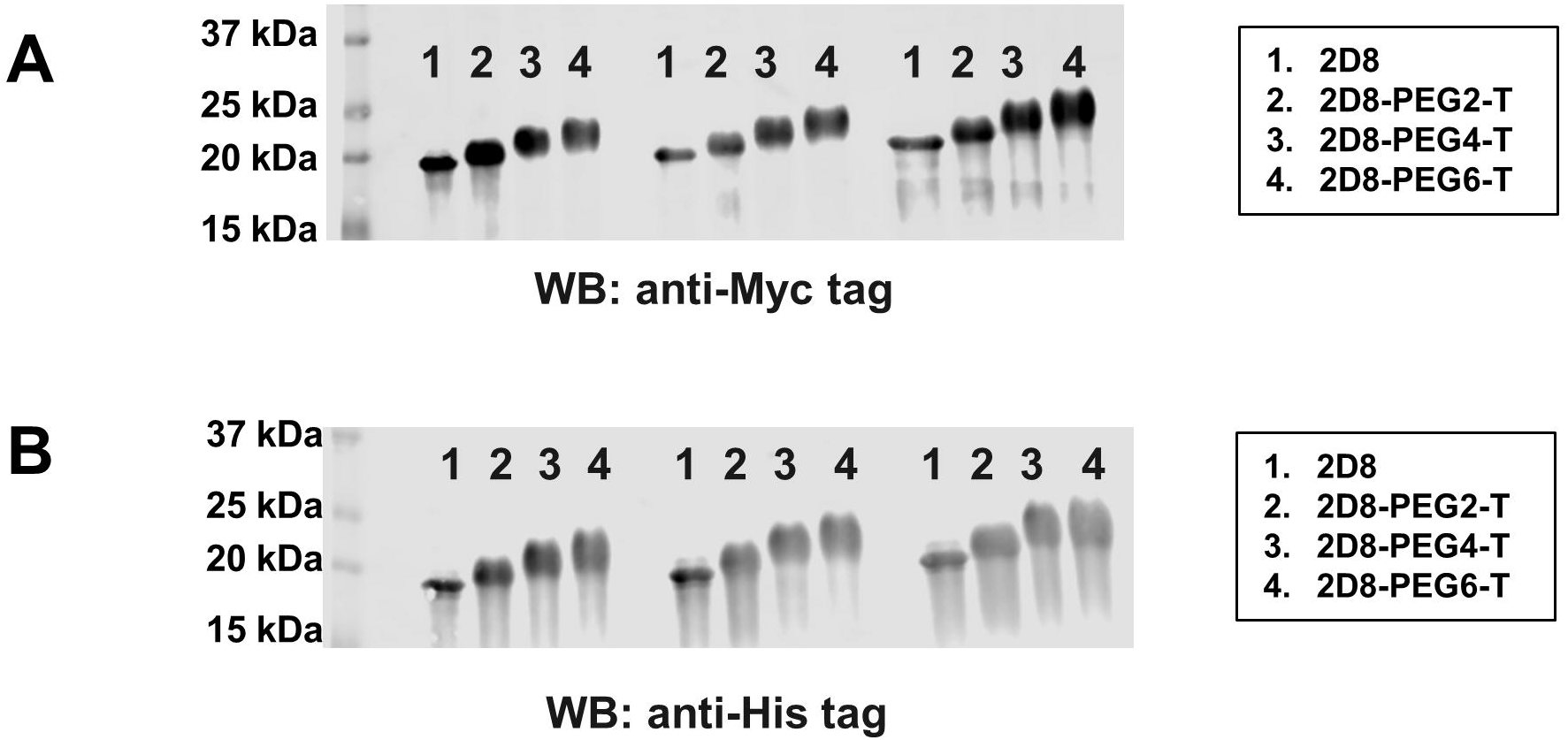
Synthesis of single-domain antibody (sdAb)-based protein degrader. sdAb 2D8 was conjugated to thalidomide (an E3 ligase ligand) using three different lengths of linkers [– (PEG)n–, n=2, 4, 6], that were attached to the lysine residues of the sdAb. Both unmodified sdAb 2D8 and the three modified sdAbs contained Myc and His tags at the N-terminal. Western blots show the detection of the sdAbs by anti-Myc antibody (A) and anti-His antibody (B).

**Supplemental Figure 2:**
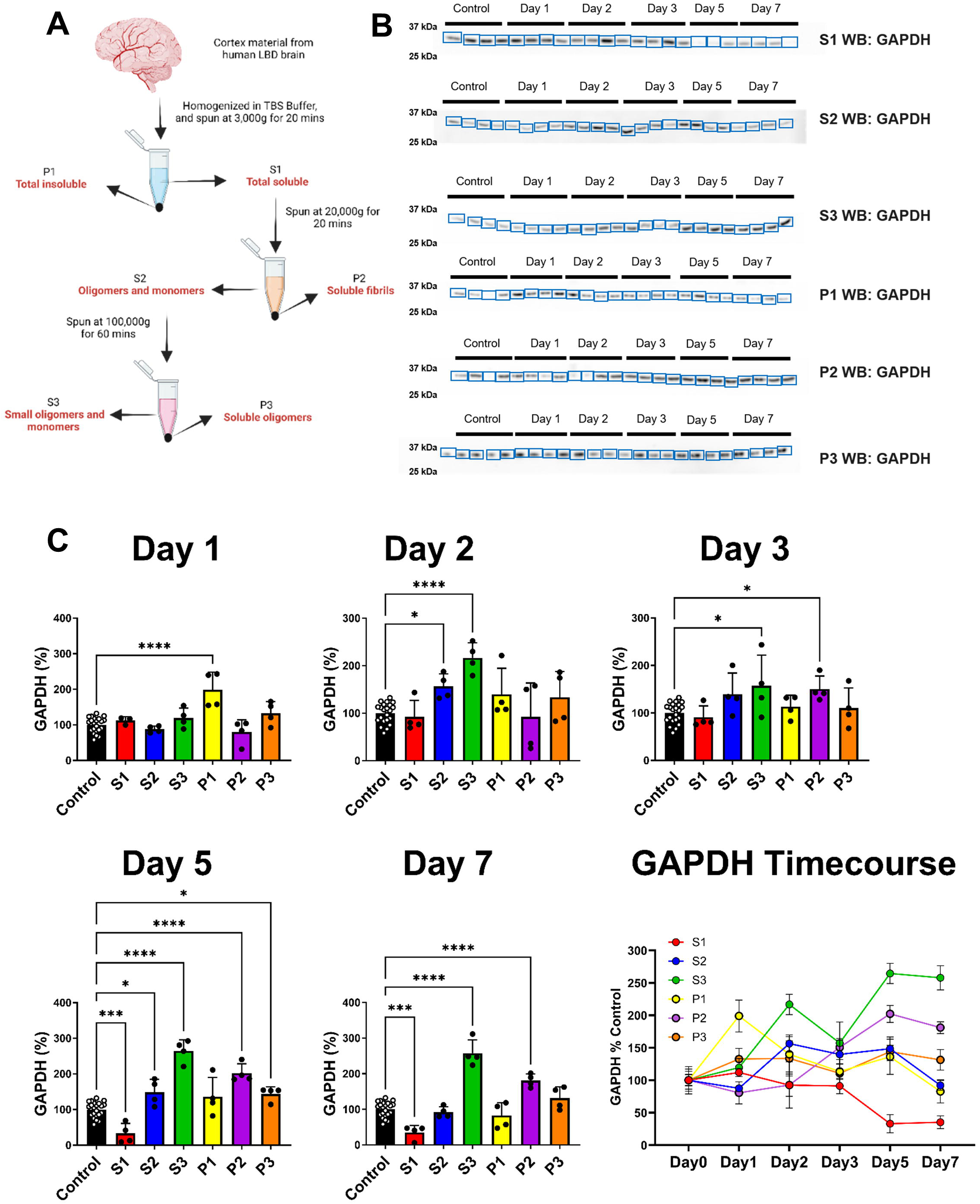
Assessment of toxicity of human Lewy body dementia (LBD) brain fractions. (A) Schematic protocol illustrating the preparation of different fraction of α-syn from LBD brain. (B-C) Immunoblots illustrating GAPDH levels in treated cell lysate on different days (B). Toxicity of different α-syn fractions of LBD brain (C). Neuronal cultures were prepared from wild-type pups at day 0. Cells were incubated for 1, 2, 3, 5 and 7 days with 10 μg/ml of each of the six brain fractions. Toxicity was assessed by western blotting for GAPDH. Day 1: A one-way ANOVA showed a significant overall treatment effect (p<0.0001). Cells treated with P1 fractions had significantly higher GAPDH levels compared to untreated control cells (p<0.0001). Day 2: A one-way ANOVA showed a significant overall treatment effect (p<0.0001). Cells treated with either S2 (p=0.0375) and S3 (p<0.0001) fractions had significantly higher GAPDH levels compared to untreated control cells. Day 3: A one-way ANOVA showed a significant overall treatment effect (p=0.0054). Cells treated with either S3 (p=0.0108) and P2 (p=0.0304) fractions had significantly higher GAPDH levels compared to untreated control cells. Day 5: A one-way ANOVA showed a significant overall treatment effect (p<0.0001). The S1 fraction produced significant toxicity when compared to untreated control cells (p=0.0005). Cells treated with either the S2, S3, P2 and P3 fractions had significantly higher GAPDH levels compared to untreated control cells. (p=0.0169, p<0.0001, p<0.0001, p=0.036). **, ***, ****: p<0.01, 0.001, 0.0001 (Tukey’s post-hoc test).

**Supplemental Figure 3:**
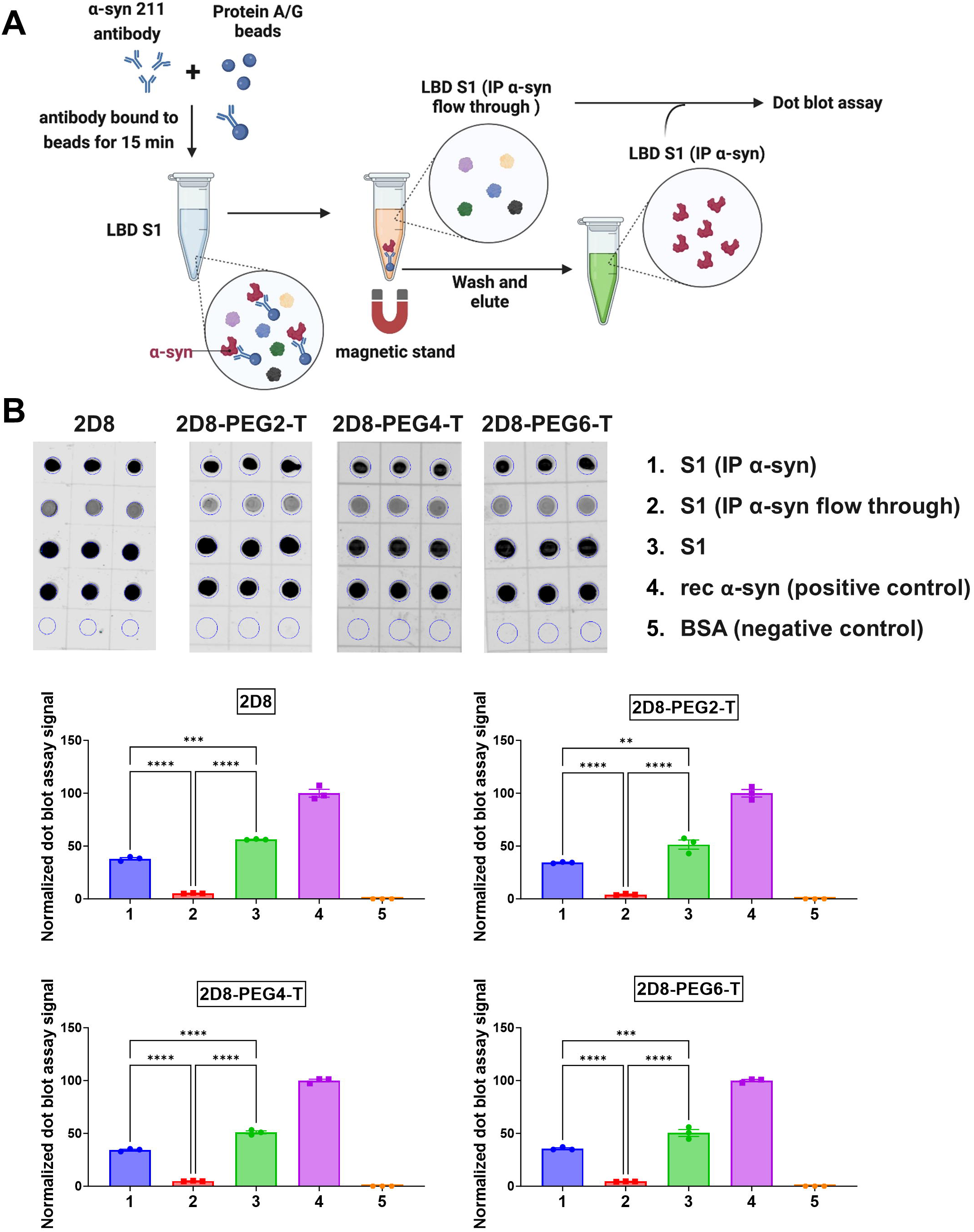
Immunoprecipitation (IP) depletion of *α*-syn from LBD S1-fraction and dot blot assay. (A) Schematic protocol for IP and dot blot assays. The experimental details are described in the Materials and Methods section. (B) Dot blot image and quantification. Positive control: rec α-syn; Negative control: bovine serum albumin (BSA). Dot blot analysis revealed significant differences in normalized signals between the groups (p<0.0001 for 2D8, 2D8-PEG2-T, 2D8-PEG4-T, and 2D8-PEG6-T; one-way ANOVA). Both 2D8 and modified 2D8 showed reduced binding after α-syn depletion, with binding decreasing by 90% (p<0.0001, 2D8), 92% (p<0.0001, 2D8-PEG2-T), 90% (p<0.0001, 2D8-PEG4-T), and 91% (p<0.0001, 2D8-PEG6-T) compared to S1, respectively. Notably, stronger binding of S1 (IP α-syn) was observed, with binding increasing by 86% (p<0.0001, 2D8), 88% (p<0.0001, 2D8-PEG2-T), 86% (p<0.0001, 2D8-PEG4-T), and 88% (p<0.0001, 2D8-PEG6-T) compared to S1 (IP α-syn flow through), respectively.

**Supplemental Figure 4:**
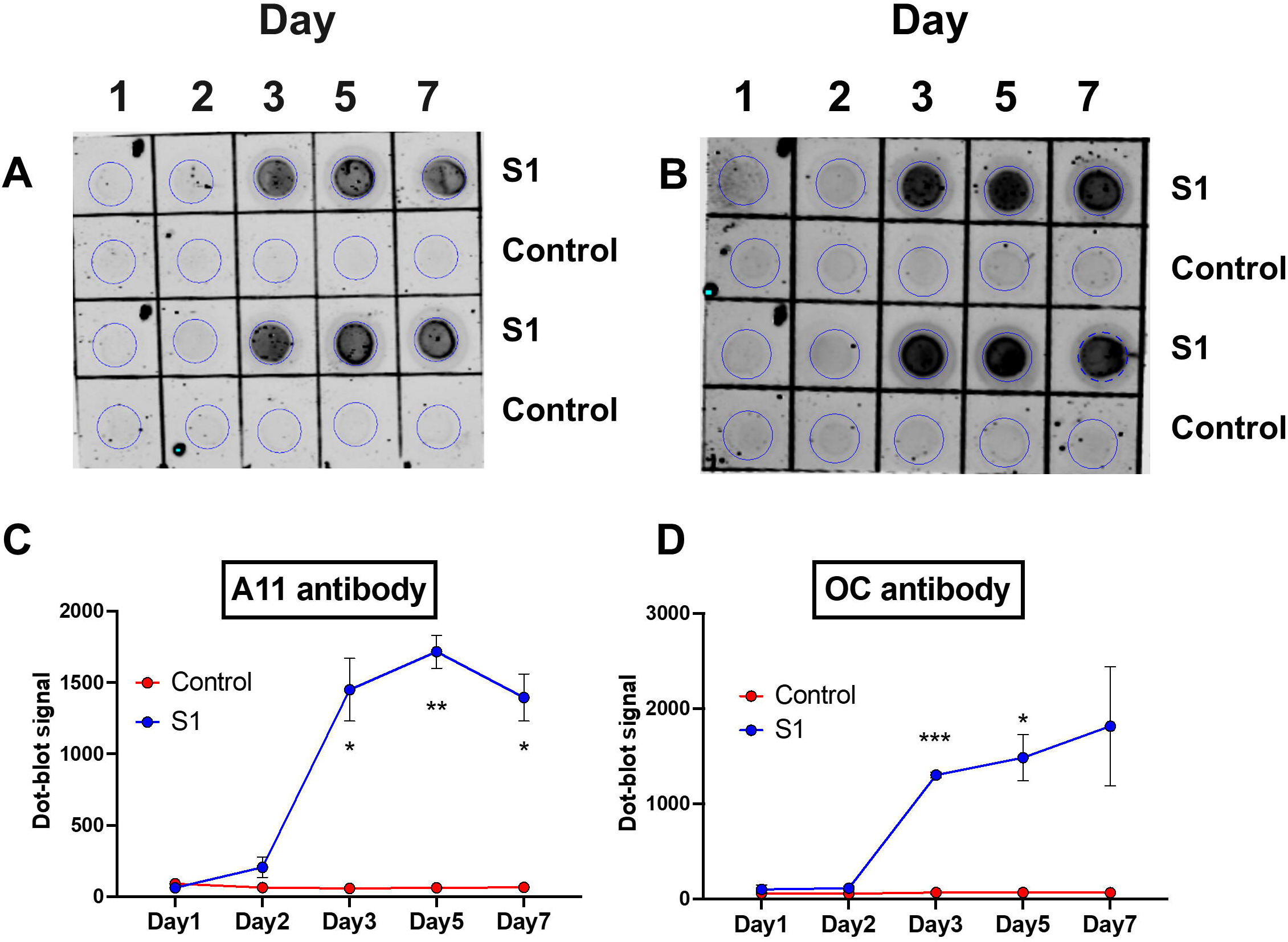
*α*-Syn seeding experiment. Neuronal cultures were established from M83 mouse pups on day 0. Cells were treated with 10 µg/ml of LBD S1 fraction for 1, 2, 3, 5, and 7 days. The α-syn seeding capacity was evaluated using dot blot assays with conformational antibodies A11 (A) and OC (B). Quantitative analysis of the dot blot signals revealed significantly increased levels of α-syn aggregates in the presence of the LBD brain soluble S1 fraction (A11 antibody (C): p=0.0241 (day 3), p=0.0048 (day 5), p=0.0151 (day 7); OC antibody (D): p=0.0007 (day 3), p=0.0286 (day 5); unpaired t-test).

**Supplemental Figure 5:**
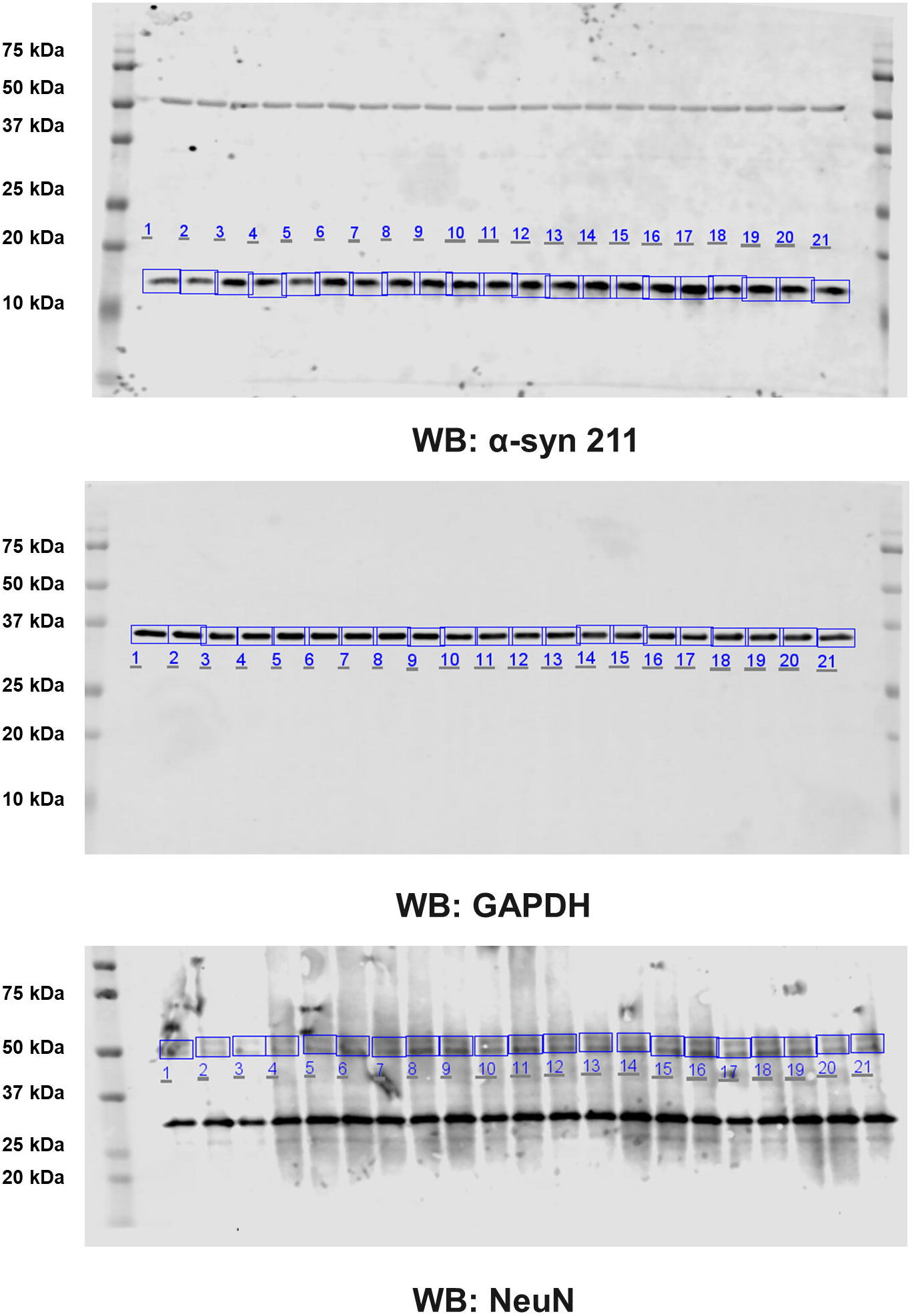
Complete western blots and bands quantified in Fig. 3.

**Supplemental Figure 6:**
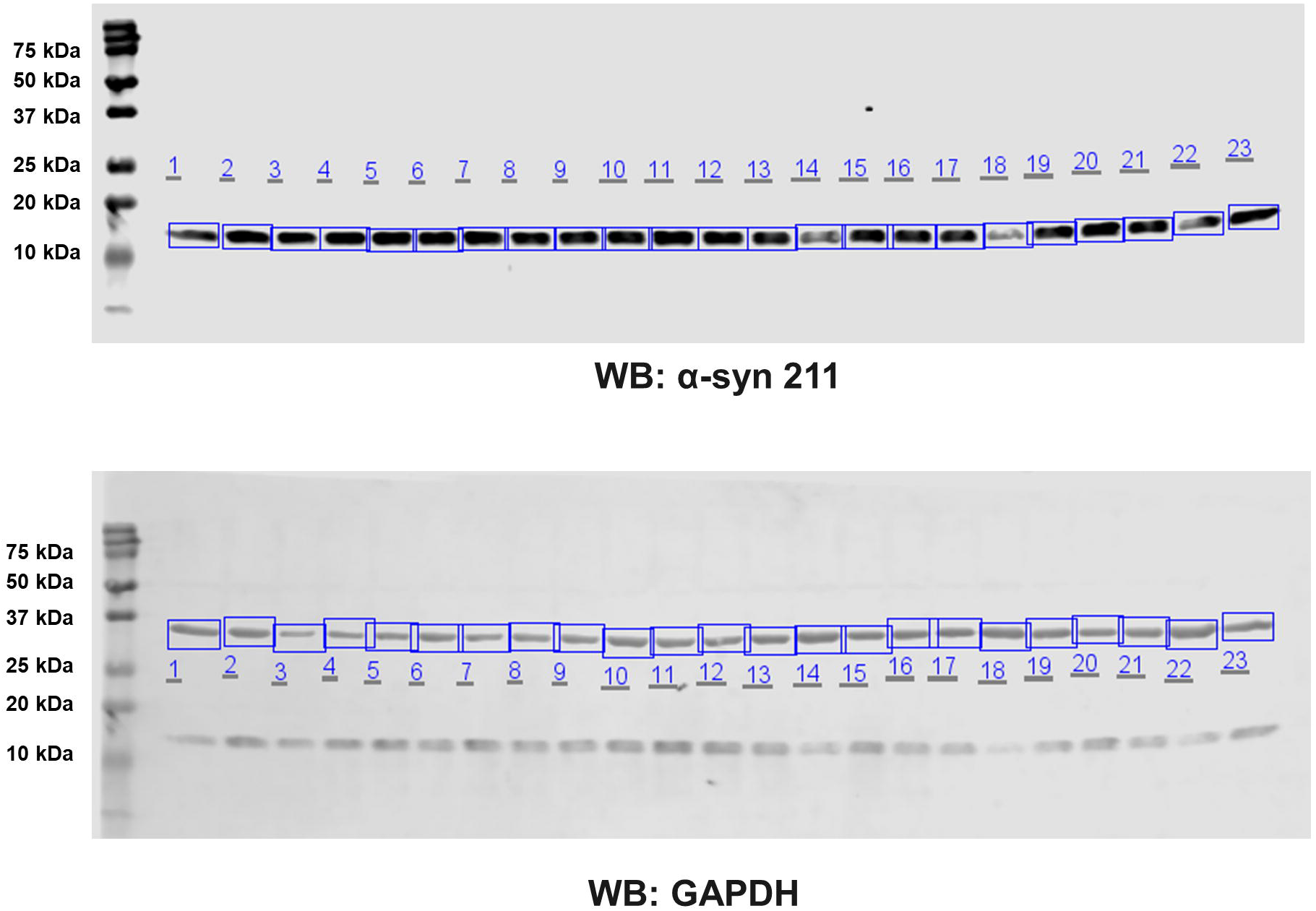
Complete western blots and bands quantified in Fig. 4.

**Supplemental Figure 7:**
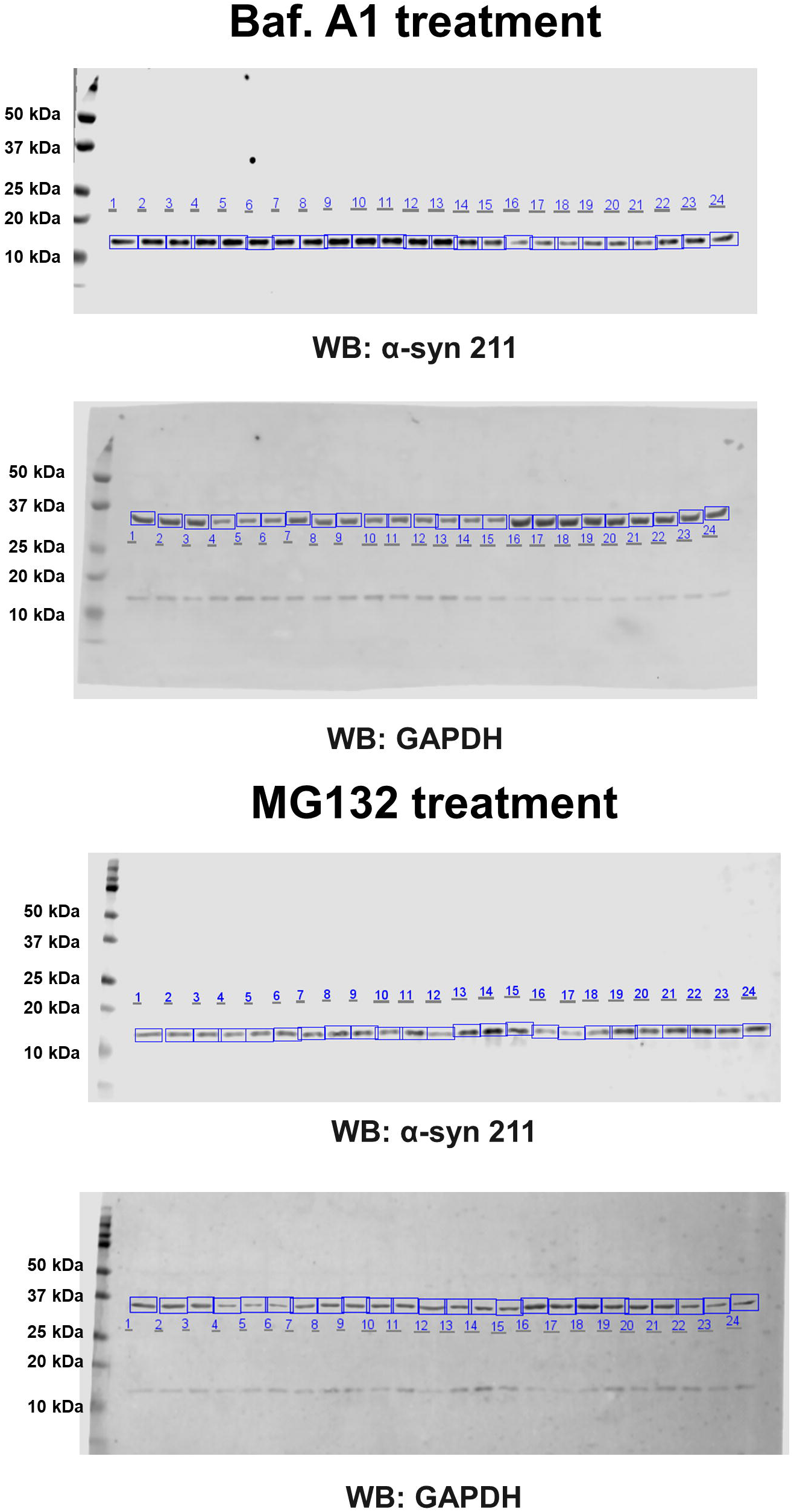
Complete western blots and bands quantified in Fig. 5.

**Supplemental Figure 8:**
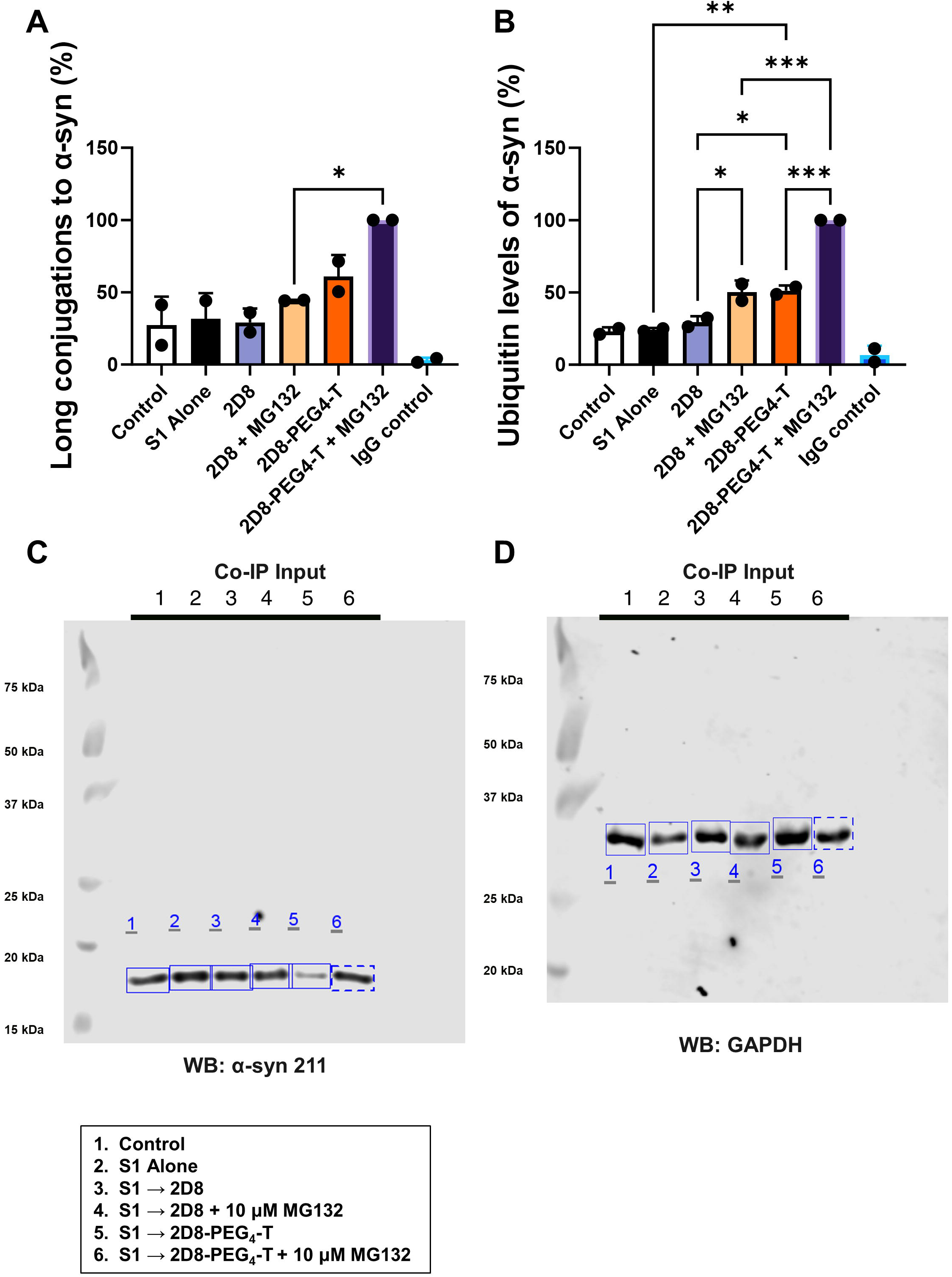
(A-B) Quantitation for the western bands presented in Figure 6 (C-D). **(A)** Long conjugations to α-syn: Significant differences were observed between groups in long conjugations to α-syn (Figure 6C (i), one-way ANOVA, p=0.0017). The modified sdAb 2D8-PEG4-T with the proteasome inhibitor MG132 resulted in a 56% increase in long conjugations to α-syn compared to untreated sdAb 2D8 with MG132 (p=0.0245). **(B)** Ubiquitin levels: Significant group differences were observed in ubiquitin levels (Figure 6D (i), one-way ANOVA, p<0.0001). The modified sdAb 2D8-PEG4-T significantly elevated ubiquitin levels by 53% (p=0.0064) and 43% (p=0.0208) compared to S1 alone and sdAb 2D8, respectively. Treatment with the proteasome inhibitor MG132 notably increased ubiquitin levels by 42% (p=0.027, sdAb 2D8 vs sdAb 2D8 + MG132) and 49% (p=0.0002, sdAb 2D8-PEG4-T vs sdAb 2D8-PEG4-T + MG132) for both unmodified and modified sdAb 2D8 groups. Furthermore, modified sdAb 2D8-PEG4-T combined with MG132 led to a 50% increase in ubiquitin levels compared to unmodified sdAb 2D8 with MG132 (p=0.0001). **(C-D)** Complete Western Blots in Figure 6E.

**Supplemental Figure 9:**
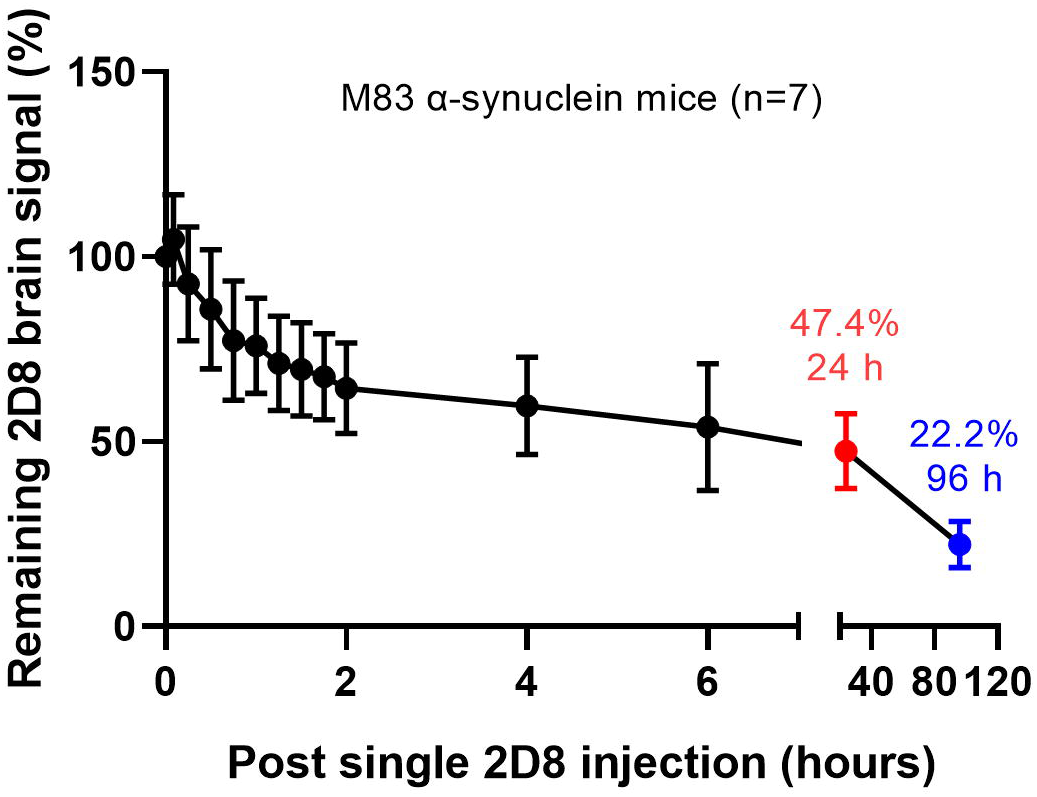
Brain-signal declines over days post single intravenous injection of near-infrared-labeled sdAb 2D8 (10 mg/kg).

**Supplemental Figure 10:**
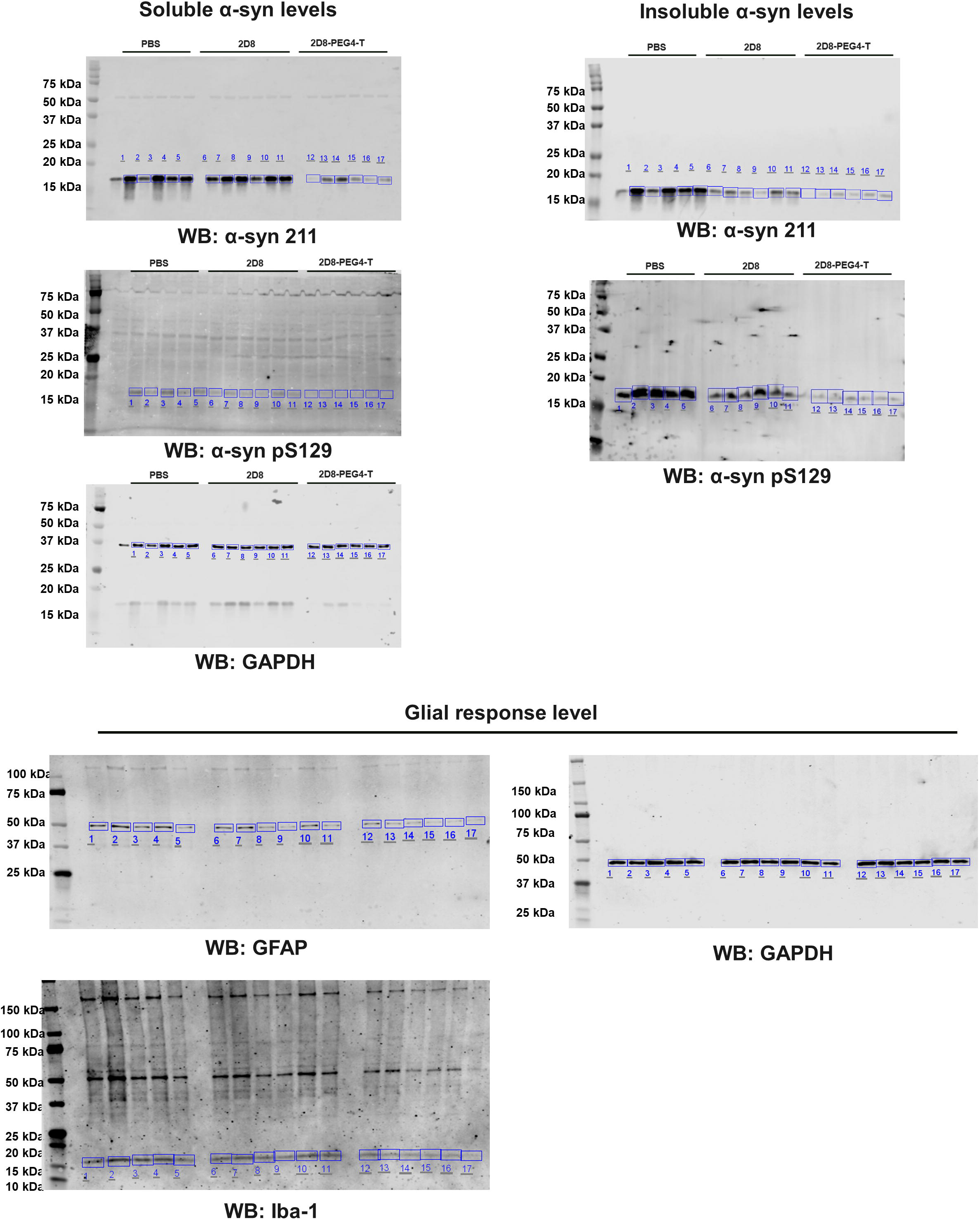
Complete western blots and bands quantified in Fig. 8-9. The single band without a box in soluble α-syn 211 and GAPDH as well as insoluble α-syn 211 is an animal that was excluded from all analyses because of a very high IVIS signal throughout the body that appeared to be non-specific.

